# A Transient High-dimensional Geometry Affords Stable Conjunctive Subspaces for Efficient Action Selection

**DOI:** 10.1101/2023.06.09.544428

**Authors:** Atsushi Kikumoto, Apoorva Bhandari, Kazuhisa Shibata, David Badre

## Abstract

Flexible action selection requires cognitive control mechanisms capable of mapping the same inputs to different output actions depending on the context. From a neural state-space perspective, this requires a control representation that separates similar input neural states by context. Additionally, for action selection to be robust and time-invariant, information must be stable in time, enabling efficient readout. Here, using EEG decoding methods, we investigate how the geometry and dynamics of control representations constrain flexible action selection in the human brain. Participants performed a context-dependent action selection task. A forced response procedure probed action selection different states in neural trajectories. The result shows that before successful responses, there is a transient expansion of representational dimensionality that separated conjunctive subspaces. Further, the dynamics stabilizes in the same time window, with entry into this stable, high-dimensional state predictive of individual trial performance. These results establish the neural geometry and dynamics the human brain needs for flexible control over behavior.

## Introduction

Integration of our abstract knowledge, goals, contexts, or internal states to select pathways for specific actions is at the basis of intelligent, adaptive behavior. For example, most of the time, we might respond to notifications of a text thread from a friend immediately. However, on occasion, the same inputs might be routed to different actions (e.g., pressing mute) depending on the prevailing context (e.g., when in a meeting with a colleague). Therefore, as this case illustrates, flexible behavior depends on our capacity to map similar or even identical inputs to many alternative and distinct actions, as the situation and our internal goals demand. This capacity is termed cognitive control. A neural representation that enables this type of context-sensitive, nonlinear mapping is a control representation (Badre et al., 2021; J. D. Cohen, 2017; E. K. Miller & Cohen, 2001). Thus, understanding the nature of control representations is fundamental for constraining theory and advancing our understanding of adaptive and flexible action selection.

Encoding of control representations is constrained by the dynamic geometrical properties of neural trajectories that unfold nonlinearly during task execution. To achieve reliable and flexible context-sensitivity for cognitive control, one critical challenge for neural systems is to assemble control representations that maintain selectivity, separability, and stability of task-critical information in a large and dynamic neural state space (Duncker & Sahani, 2021; Gao et al., 2017; Langdon et al., 2023; Stokes et al., 2013; Vyas et al., 2020). Specifically, for action selection, a control representation should (i) be selective to task-critical information about the input, output or goal, (ii) encode highly similar neural input states in a format that separates task-critical dimensions for different contexts, and (iii) provide temporally stable readout subspaces to downstream neurons that can bridge variable gaps in time. In other words, control representations with robust context-sensitivity must achieve dynamic geometrical properties that enhance separability while preserving the stability of neural trajectories for the selective extraction of task-critical information.

The separability of a neural population encoding control representations can be modulated by its geometric organization. For instance, representational dimensionality–the minimum number of axes needed to account for the variability of neural patterns that span across the task input conditions– is known to balance a computational tradeoff between separability and generalizability of representations in terms of available readouts (Ahlheim & Love, 2018; Barak et al., 2013; Fusi et al., 2016; Jazayeri & Ostojic, 2021a; Johnston et al., 2020). Specifically, a higher-dimensional representational geometry allows the system to encode even similar inputs into more separable or orthogonal neural patterns, thus reducing the overlap of the patterns (Figure 1A). Thus, representations encoded in a high-dimensional format could be beneficial for cognitive control, in that changing contextual contingencies often requires distinguishing highly correlated inputs (Badre et al., 2021; Rigotti et al, 2010).

**Figure 1.**
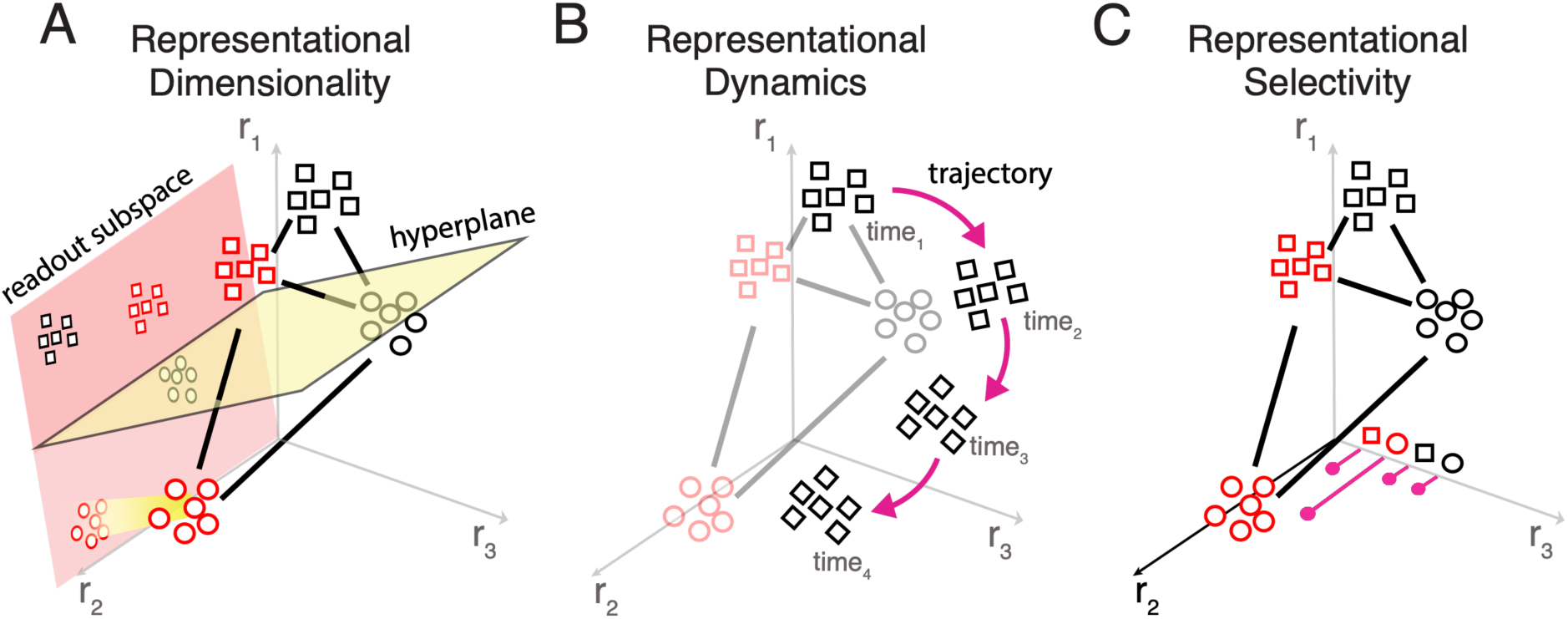
Representational dimensionality, dynamics and selectivity. Schematic illustration of representational dimensionality, dynamics, and selectivity of the conjunctive control representations. Each panel plots the response of a toy population of three units to input conditions varying in shape (square or circle) and color (red or black). Axes represent the firing rates of single units, collectively defining a neural activity space for the population. Each point within this activity space represents the population response for a given input (identified by colored shapes). Distance between points reflects how distinct responses are, and the jittered cloud of points reflect the trial-by-trial variability in responses to a given input. (A) The geometric format of the population neural responses is defined by response patterns arranged in 3 dimensions. A linear readout is implemented by a decision hyperplane (yellow) that divides the readout subspace into different classes (e.g., different shapes of inputs). The high-dimensional representation (traced by solid black lines), where no cluster of responses are aligned to each other, allows a wider variety of input conditions to be linearly separable. In addition, responses projected to a readout subspace defined by a linear hyperplane tend to dissociate input conditions (e.g., red-square and black-square) that are on the same side of the decision boundary, increasing the separability of neural responses overall. (B) Due to the time-varying nature of neural activity, the neural state space is changing and reshaping its underlying geometry over time. The dynamic neural trajectories potentially require changes in the weights for optimal linear hyperplanes for downstream readouts. (C) Units within the population show a heterogeneous turning profile or nonlinear mixed selectivity. To illustrate, the tuning profile for one unit (r_2_) is plotted along the corresponding axis. The bars plot activity of r_2_ to each of the four conditions of input and depict a non-linear mixed selective pattern. Note that similar geometric properties are expected at the level of a single unit (i.e., mixed-selectivity) or population of neurons (i.e. integrative subspaces). The event-file representations could be conceptualized as one form of mixed selectivity where a unit or groups of units are exclusively tuned to a specific combination of task critical factors more than other pairs.

However, achieving stability for temporal invariance of readout, while enhancing separability for context-sensitivity via high-dimensional geometry, often poses a computational challenge. For example, recurrent neural networks that use chaotic dynamics or reservoir computing can achieve exceptionally high context-sensitivity, yet they are temporally unstable (Buonomano & Maass, 2009; Cueva et al., 2020; Enel et al., 2016; Farrell et al., 2022; Maass et al., 2002). Such fully dynamic coding makes reliable readout of task-critical information difficult because the selectivity and geometry of neural populations could change over time (Murray et al., 2017). This makes computed solutions inefficient and not generalizable to other event timings (Rigotti et al, 2010).

The problem of time-invariance can be mitigated by learning stable, temporally low-dimensional, attractor-based dynamics where trajectories span subspaces or manifolds that offer a fixed, common set of weights for readout (Druckmann & Chklovskii, 2012; P. Miller, 2016). Indeed, neural computations of task-critical information often involve partially dynamic yet structured neural trajectories (Driscoll et al., 2022; Enel et al., 2020; Kozachkov et al., 2020; Mante et al., 2013; Okazawa et al., 2021; Schuessler et al., 2020; Spaak et al., 2017), reaching stable subspaces to represent information despite time-varying dynamics in the overall neural state space (Bouchacourt & Buschman, 2019; Murray et al., 2017; Panichello & Buschman, 2021; Parthasarathy et al., 2019; Tang et al., 2020). Thus, the fundamental question is what kind of stable subspaces could provide context-sensitivity with enhanced separability that benefits cognitive control problems.

A possible solution to enhance separability while maintaining stability for robust cognitive control is to selectively encode stable conjunctive subspaces that integrate task-critical information within a high-dimensional representational geometry. The expansion of dimensionality affords the integration of low-dimensional task features such as the abstract context and the settings for inputs and outputs. This solves the context-sensitivity problem and enhances separability by flexibly encoding dissociable context-dependent mappings. Further, achieving stable dynamics (e.g., fixed points) within conjunctive subspaces provides the temporal invariance that facilitates more stable readout even while neural trajectories are highly dynamic in the overall neural state space.

In line with this hypothesis, several theories and computational models of cognitive control suggest that robust selection of context-dependent actions requires encoding and retrieval of conjunctive representations, which bind key task features such as goals, rules, stimuli, responses, and outcomes (Frings et al., 2020; Hommel et al., 2001; Verguts & Notenaert, 2009; Ito et al., 2022; Logan, 1989; Schumacher & Hazeltine, 2016; Verbeke & Verguts, 2022). Supporting these theories, a line of studies decoding EEG signals in humans during performance of response selection tasks has found consistent evidence of conjunctive representations supporting context-dependent action selection (Kikumoto, Mayr, et al., 2022; Kikumoto, Sameshima, et al., 2022; Kikumoto & Mayr, 2020; Rangel et al., 2023; Takacs et al., 2020).

Notably, conjunctive subspaces can be easily implemented as a neural population code that exhibits nonlinear mixed-selectivity (Asaad et al., 2000; Bouchacourt & Buschman, 2019; Dang et al., 2021; Fusi et al., 2016; Kaufman et al., 2022; Kira et al., 2023; Lindsay et al., 2017; Rigotti et al, 2010; Rigotti et al., 2013); Figure 1C) or larger scale task-contingent interactions of neural assemblies that effectively expand representational dimensionality of population codes (Buschman et al., 2012; Ito et al., 2022; Semedo et al., 2019; van den Brink et al., 2022). Within such a high-dimensional, expressive geometry, readouts of task-critical information, including conjunctions, are expected to be enhanced.

Finally, neural network models with nonlinear recurrent dynamics solving context-dependent tasks show stable dynamics that implement subspaces or manifolds across task-critical dimensions, increasing the adaptability of learned representations (Cueva et al., 2020; Druckmann & Chklovskii, 2012; Egger et al., 2019; Pollock & Jazayeri, 2020; Rajalingham et al., 2022; Shenoy et al., 2013). Therefore, if conjunctive representations are encoded in stable subspaces on the fly, cognitive control could operate by directly implementing readouts of context-dependent input-output mappings that are minimally required by the task, recruiting stable yet separable neural patterns (Figure 1 A, B, and C). However, no direct links between such dynamic geometric properties of control representations and controlled behavior has been established in humans.

In the current study, we sought to characterize the nature of control representations in terms of their geometric properties and dynamics during context-dependent action selection in humans. Specifically, we hypothesized that temporally stable conjunctive subspaces that are expressed in a high-dimensional representational geometry provide enhanced separability that facilitates the readout of specific controlled actions. Conversely, responses made before the computation of a stable conjunctive state within a high-dimensional geometry cannot take full advantage of greater separability and will be more likely to produce errors.

To test this hypothesis, we combined the rule-based action selection task with a psychophysical response-deadline procedure and several time-resolved decoding techniques for EEG (Figure 2; see *Methods* for detail). After varying time intervals, participants were cued to make context-dependent actions immediately. This procedure compelled responses based on the current state of neural representations, thereby capturing the readout of task-critical information at distinct stages during the progression of representational neural trajectories. This allowed us to evaluate the dimensionality, dynamics, and selectivity of task representations governing behavior (Figure 1).

**Figure 2.**
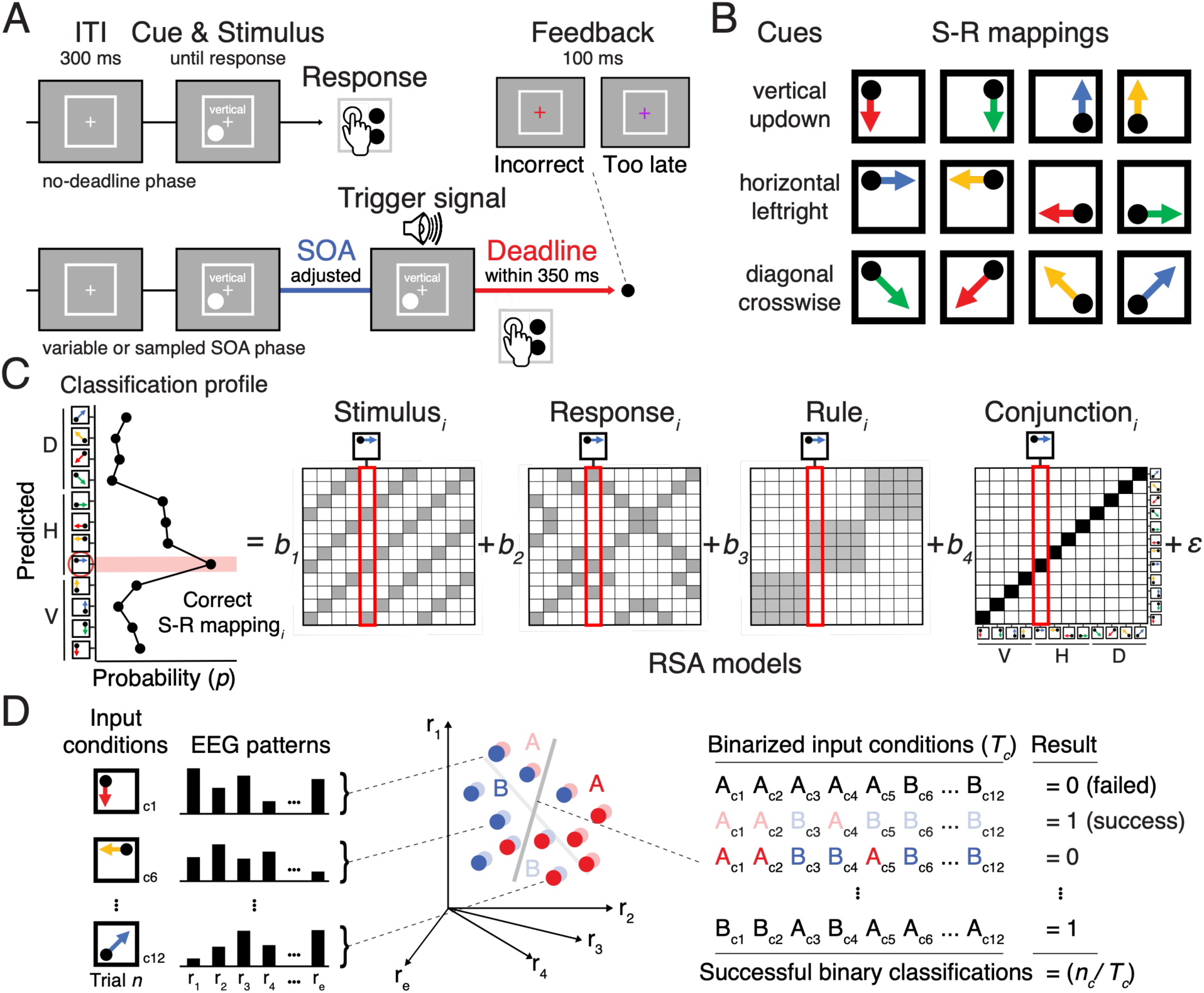
Task design and the procedure of decoding analyses. (A) Sequence of trial events in the rule-selection task with the response deadline. In the variable or sampled SOA phase, an audio trigger signal indicated the start of a response deadline time window. (B) Spatial translation of different rules (rows) mapping different stimuli (columns) to responses (arrows), yielding 12 independent conjunctions. (C) Schematic of the time-resolved representational similarity analysis. For each sample time (*t*), a scalp-distributed pattern of EEG was used to decode the specific rule/stimulus/response configuration of a required action. The decoder produced sets of classification probabilities for each of the possible action constellations. The profile of classification probabilities reflects the similarity structure of the underlying representations, where action constellations with shared features are more likely to be confused. For each trial and timepoint, the classification probabilities were regressed onto model vectors as predictors that reflect the different, possible representations. In each model matrix, the shading of squares indicates the theoretically predicted classification probabilities (darker shading means higher probabilities) in all possible pairs of constellations. The coefficients associated with each predictor (i.e., *t-*values) reflect the unique variance explained by each of the constituent features and their conjunction. (D) Schematic of the time-resolved binary classification method used to estimate the representational dimensionality (Rigotti et al., 2013). For each time point (*t*), a pattern of EEG associated with unique action constellations or input conditions (c_1-12_) take a position in the multidimensional neural space spanning in r_1-e_ dimensions. By assigning new binary class labels (e.g., A or B) to the input conditions, we can generate arbitrary binary groupings given the task conditions. Because the higher dimensional geometry of neural responses generally affords more arbitrary linear separations by being more expressive, the count of successfully implementable binary classifications of newly defined groupings scales with the representational dimensionality. To adapt the method to EEG by lowering the cutoff threshold for classification, we used an exclusive cutoff method. In this method, classifications must exceed the cutoff threshold in all the different input conditions assigned to each trial, as opposed to the averaged performance across c_1-12_, to be marked as success.

To preview our results, we discovered that successful action selection recruits a transient expansion of representational dimensionality, whose peak is coincident with temporal stability in a conjunctive subspace established just prior to response execution. Collapsed dimensionality led to incorrect and/or slow responses. In addition, earlier development of temporally stable conjunctive subspaces predicted unique variance of the quality of trial-to-trial action selection. These results provide evidence that highly separable yet temporally stable dynamics in the coding of conjunctive control representations are important for efficient goal-contingent action selection.

Participants performed this rule-based selection task in three task phases in order: the no-deadline phase, the variable stimulus-onset asynchrony (SOA) phase, and the sampled SOA phase. In the no-deadline phase, after 15 practice blocks, participants performed the rule-based action selection task and were only instructed to respond as accurately and quickly as possible, balancing both equally. The no-deadline phase provided reference RTs, which were used to adjust the timing of SOA intervals for subsequent phases as described in the Estimation of SOA functions (Figure S1A).

## Results

### Behavior in the no-deadline, variable SOA and sampled SOA task

During the no-deadline task phase, participants learned to perform the task rapidly and accurately (*M* = 620 ms, *SE* = 69 ms for RTs and *M* = .07, *SE* = .03 for error; Figure S1A), following ex-gaussian distributions (Figure S1B).

Next, participants performed the same task with response deadline and adaptive stimulus onset asynchrony (SOA) intervals between the onset of stimulus and the response deadline. We term this phase the variable SOA task. As a function of increasing SOA, the rate of correct responses (i.e., accurate responses that occurred within response deadline) significantly increased then plateaued (Figure 3A), *t*(1,41) = 23.31, *beta* = 5.10, 95 % CI [4.67 5.53] for linear effect, and *t*(1,41) = -17.97, *beta* = -2.35, 95 % CI [-2.60 -2.94] for quadratic effect. Focusing on accurate and within-deadline responses, participants responded faster as SOA increased (*M* = 248 ms, *SD* = 17.76 ms). Both complete omission of responses (*M* = 1.83 %, *SD* = 1.58 %) and premature responses emitted before the trigger signal (*M* = 2.70 %, *SD* = 2.70 %) were rare. Further, as the SOA intervals increased, the omission of responses (*M* = 1.83 %, *SD* = 1.58%) occurred significantly less, and premature responses occurred more (Supplementary table 1).

**Figure 3.**
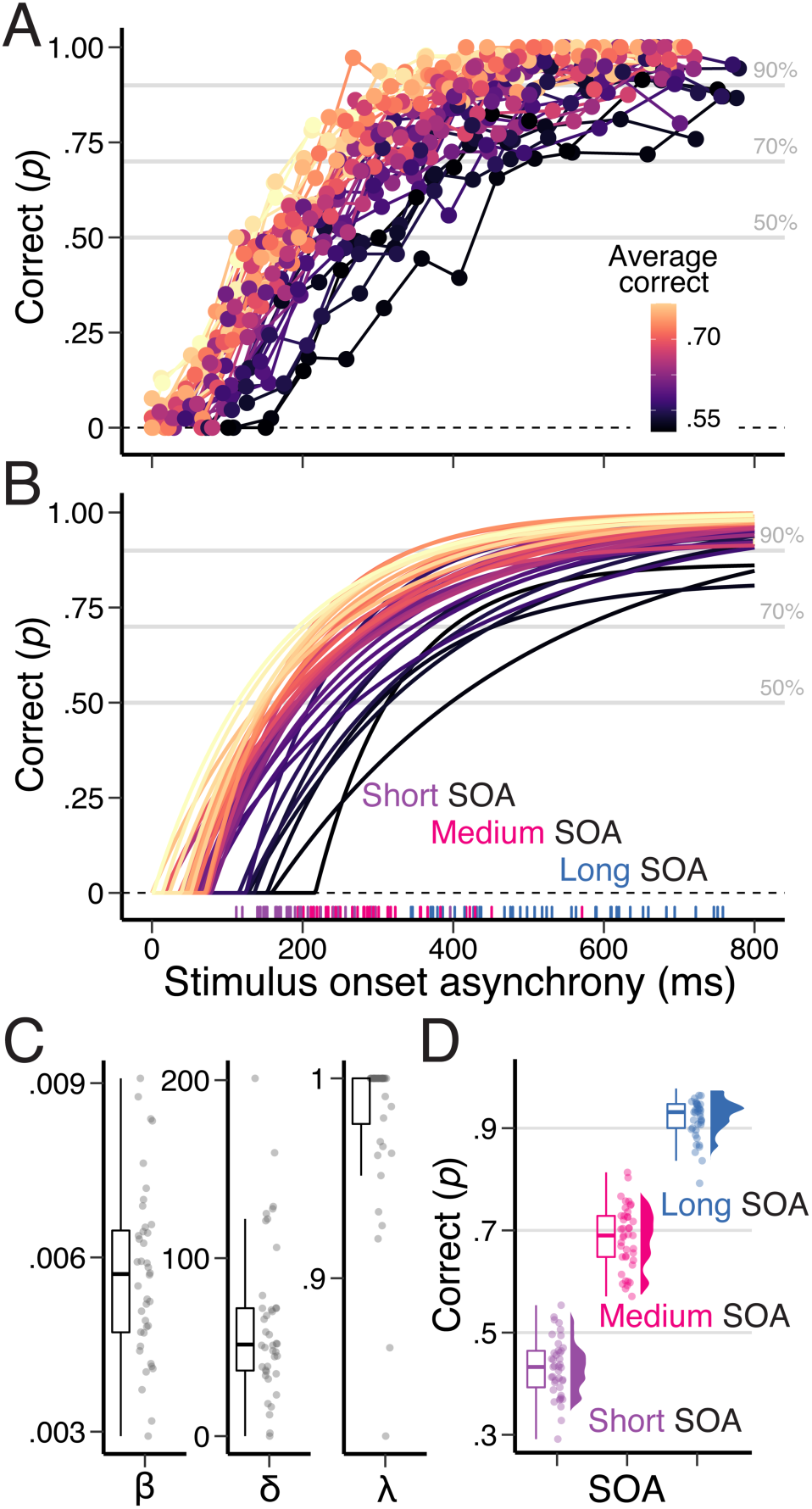
Psychophysical curve of on-time response accuracies via SOA manipulation. (A) Empirically observed changes in the probability of correct and on-time responses as a function of the subjects-specific SOA intervals. Each line plots an individual subject. (B) Predictions by the model for each subject of an exponential function approaching to the limit using the subjects-specific estimates of parameters. (C) Distributions of the parameters of the model with an exponential function to the limit: β is the rate parameter, which indexes the speed at which accuracy grows to asymptote; δ is the intercept reflecting the discrete point when accuracy departs from 0, and λ is the asymptotic accuracy level reflecting the overall probability of successful responses. (D) Empirically observed probability of correct responses during sampled SOA phase where we simultaneously measured EEG. The subjects-specific SOA intervals estimated from earlier phases were used to force expected number of errors for short SOA (*p* = .5), medium SOA (*p* = .75), and long SOA (*p =* .9) condition respectively. (C-D) The box plots show 25^th^, 50^th^, and 75^th^ percentile as a box and minima/maxima of whiskers correspond to smallest/largest values within 1.5 times interquartile range below/above the 25^th^/75^th^ percentile.

Inappropriate responses consisted of response selection errors or responses that were too late for the deadline, and both types of errors drove adaptive SOA adjustments. On average, 28.86% (*SD* = 7.23%) of responses occurred after the deadline. Excluding complete omission trials (i.e., no response at all) and across all SOA conditions, response commission errors occurred on 10.04% (*SD* = 6.28%) of trials within the deadline and 29.43% (*SD* = 11.29%) after the deadline (see Figure S2 for the distributions of trials of different response types).

Commission response errors significantly decreased as a function of SOA, *t*(1,41) = - 11.27, *beta* = -3.93, 95% CI [-4.62 -3.25] for linear effect, and *t*(1,41) = -.16, *beta* = -.25, 95% CI [-.34 -.29] for quadratic effect for responses that occurred within the deadline; *t*(1,41) = -2.47, *beta* = -.62, 95% CI [-1.12 -.13] for linear effect, and *t*(1,41) = 3.11, *beta* = .52, 95% CI [.19 .85] for quadratic effect for responses that occurred after the deadline. A majority of late (i.e., after deadline) responses occurred in shorter SOA conditions and occurred significantly less as participants prepared for a longer time, *t*(1,41) = -32.87, *beta* = -6.48, 95% CI [-6.87 -6.09] for linear effect, and *t*(1,41) = 11.78, *beta* = 1.42, 95% CI [1.19 1.66] for quadratic effect. Thus, longer SOAs led participants to implement on-time and accurate action selection.

To force response errors at the expected rates, we modeled observed psychometric functions from the adaptive SOA procedure with an exponential rise approaching a plateau as SOA increases (Öztekin & McElree, 2010) *see Estimation of SOA functions* for detail). This allowed us to describe changes of performance in a continuous manner (Figure 3A) and estimate individuals-specific SOA intervals that are expected to produce at specific error rates (50%, 70%, and 90%; Figure 3B and 3C).

As a result, during the sampled SOA task while EEG was recorded, correct responses were observed at calibrated rates (Figure 3D). Further, we observed a similar behavioral pattern during the EEG experiment to the variable SOA task in terms of RTs, omission errors and commission errors as a function of short, medium and long SOA (Supplementary table 1). A majority of late responses occurred again in shorter SOA conditions 43.34% (*SD* = 10.31%) for short SOA, 20.39% (*SD* = 6.98%) for medium SOA, and 3.67% (*SD* = 2.41%) for long SOA. Excluding complete omission trials, commission errors occurred within the deadline at 20.37% (*SD* = 9.90%) for short SOA, 12.21% (*SD* = 5.70%) for medium SOA, and 3.93% (*SD* = 2.57%) for long SOA. These rates increased to 29.98% (*SD* = 10.18%) for short SOA, 21.67% (*SD* = 8.66%) for medium SOA, and 18.05% (*SD* = 11.30%) for long SOA for responses that occurred after the deadline (Supplementary table 1).

Again, the commission response errors significantly decreased as a function of SOA, *t*(1,41) = -15.30, *beta* = -1.36, 95% CI [1.18 1.53] for linear effect, and *t*(1,41) = -7.50, *beta* = -.29, 95% CI [-.36 -.21] for quadratic effect for responses that occurred within the deadline; *t*(1,41) = -6.33, *beta* = -.47, 95% CI [-.61 -.32] for linear effect, and *t*(1,41) = 2.24, *beta* = .10, 95% CI [.01 .20] for quadratic effect for responses that occurred after the deadline. The effect of SOA on RT, the occurrence of late responses, and premature responses were also similar to the pattern observed in the variable SOA task (Supplementary table 1). Overall, the quality and success of action selection was strongly influenced and regulated by SOA between the onset of trial to the deadline signal.

### Task Representational Selectivity

We first characterized time-dependent changes in encoding of various task representations during controlled actions while being forced to readout task information so a response could be made within the deadline determined by the three SOAs (Figure 3D). By analyzing the representational similarity structure, we decoded the representations of the rule, stimulus, response and conjunction of a prepared action from the moment-to-moment patterns of EEG activity (Figure 2C).

Consistent with our previous reports using a similar task (Kikumoto & Mayr), we found that a conjunctive representation of the action rule, stimulus and response develop before response execution over and above the individual representations of those same constituent task features (Figure 4; see Figure S4 and Figure S5 for the stimulus-aligned and the trigger-signal aligned results). The trajectories remained identical when the number of trials for correct, incorrect, and too late trials were subsampled during decoding analyses (Figure S3). In short and medium SOA conditions, the strength of task representations except stimulus features were higher in trials that led to accurate and within-deadline responses, *t*(1,41) = 4.72, *beta* = .04, 95% CI [.05 .22] for rule, *t*(1,41) = -.68, *beta* = -.01, 95% CI [-.02 .01] for stimulus, *t*(1,41) = 5.97, *beta* = .05, 95% CI [.03 .06] for response, and *t*(1,41) = 3.10, *beta* = .02, 95% CI [.01 .04] for conjunction (see Supplementary table 2 for the stimulus-aligned and the trigger-signal aligned results). However, no significant increase in decoding strength was observed between these two SOA conditions (Supplementary table 3), suggesting that merely increasing SOA intervals (i.e., pre-deadline response selection time) does not immediately translate to improved decoding performance.

**Figure 4.**
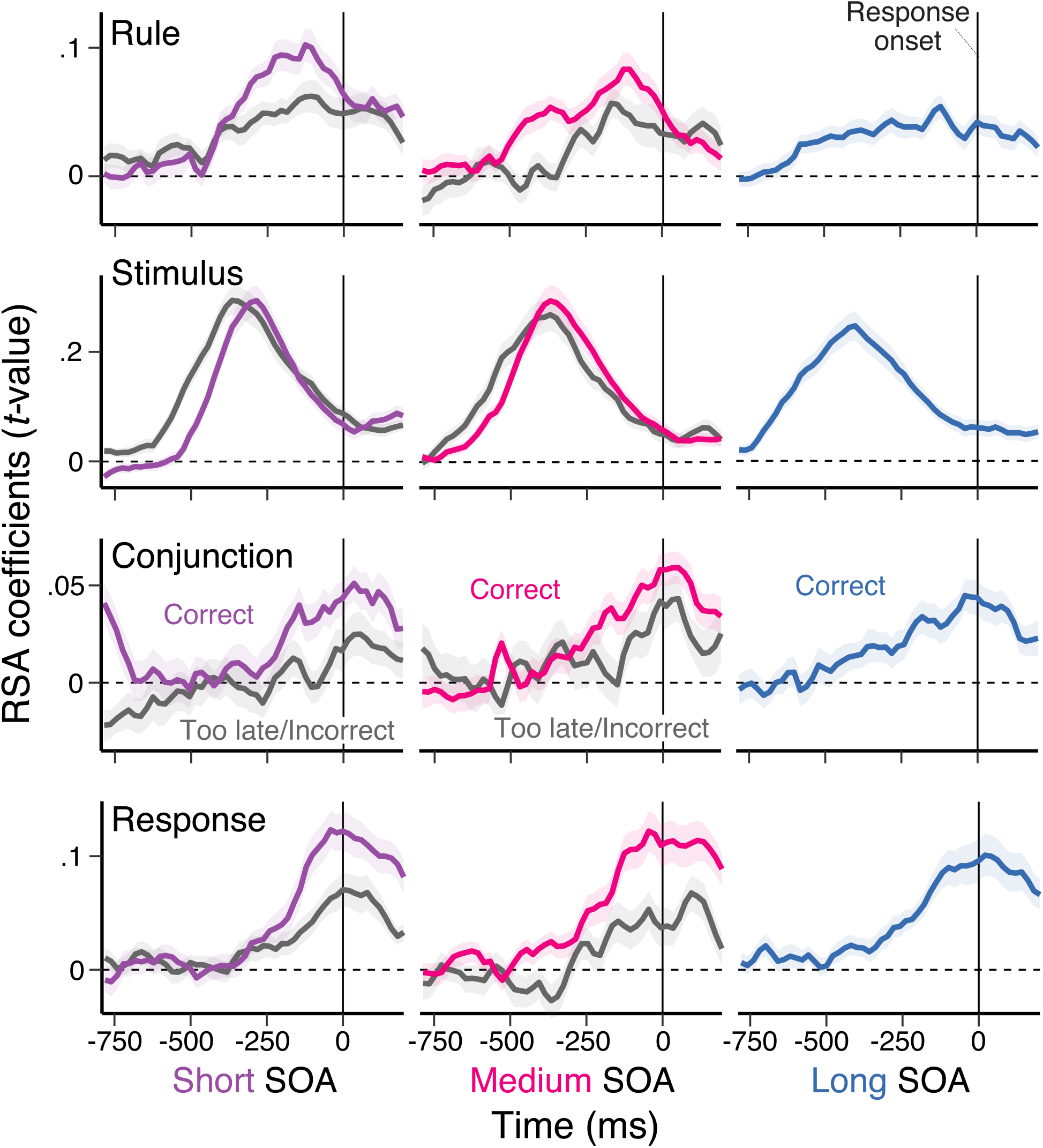
Response-aligned time-course of decoding of task representations. Average, single-trial RSA coefficients (*t*-values) associated with each of the basis set task features (rule, stimulus, and response) and their conjunction that are aligned to the onset of trial-to-trial responses. The left, middle and right column correspond to the short, medium and long SOA conditions respectively. The two lines in each panel show the average scores in trials comparing correct vs. too late or incorrect responses. The patterns were aligned to the onset of responses in each trial. Shaded regions specify the 95% within-subject standard errors.

We next related decoding strength with behavior. Higher decodability of each task representation was independently correlated with higher probability of making accurate response selections within the deadline (Figure 5 top panel) on a trial-wise basis. A similar predictive effect of response accuracy was observed in trials where responses occurred within the deadline (Figure 5 bottom panel). Notably, the effect of a rule and conjunctive representation on successful responses shift to earlier time points (-280ms before the onset of a trigger for deadline) in medium SOA trials compared to short SOA trials (Figure S7). Indeed, predictability of response failures by earlier (i.e., -400 ms prior to the trigger onset) signals encoding conjunctive representations was enhanced for longer SOAs, *t*(1,41) = 2.85, *beta* = .004, 95% CI [.01 .05] for linear effect and *t*(1,41) = 2.85, *beta* = .004, 95% CI [.01 .05] for quadratic effect, controlling for all main and interaction effects of SOAs and other task factors (see also the *Predicting Accurate Responses within Deadline Separately in Separate SOAs* in Supplementary information for detail). Further analyses revealed that responses that are too late or incorrect both decrease the strength of conjunctions to a similar degree (Figure S6), whereas the cued response representations were absent when committed responses were incorrect. Thus, more reliable encoding of the key task variables was critical for response selection under time pressure. These observations replicate and extend prior findings that the independent contribution of conjunctive representations that is unique for the prepared action.

**Figure 5.**
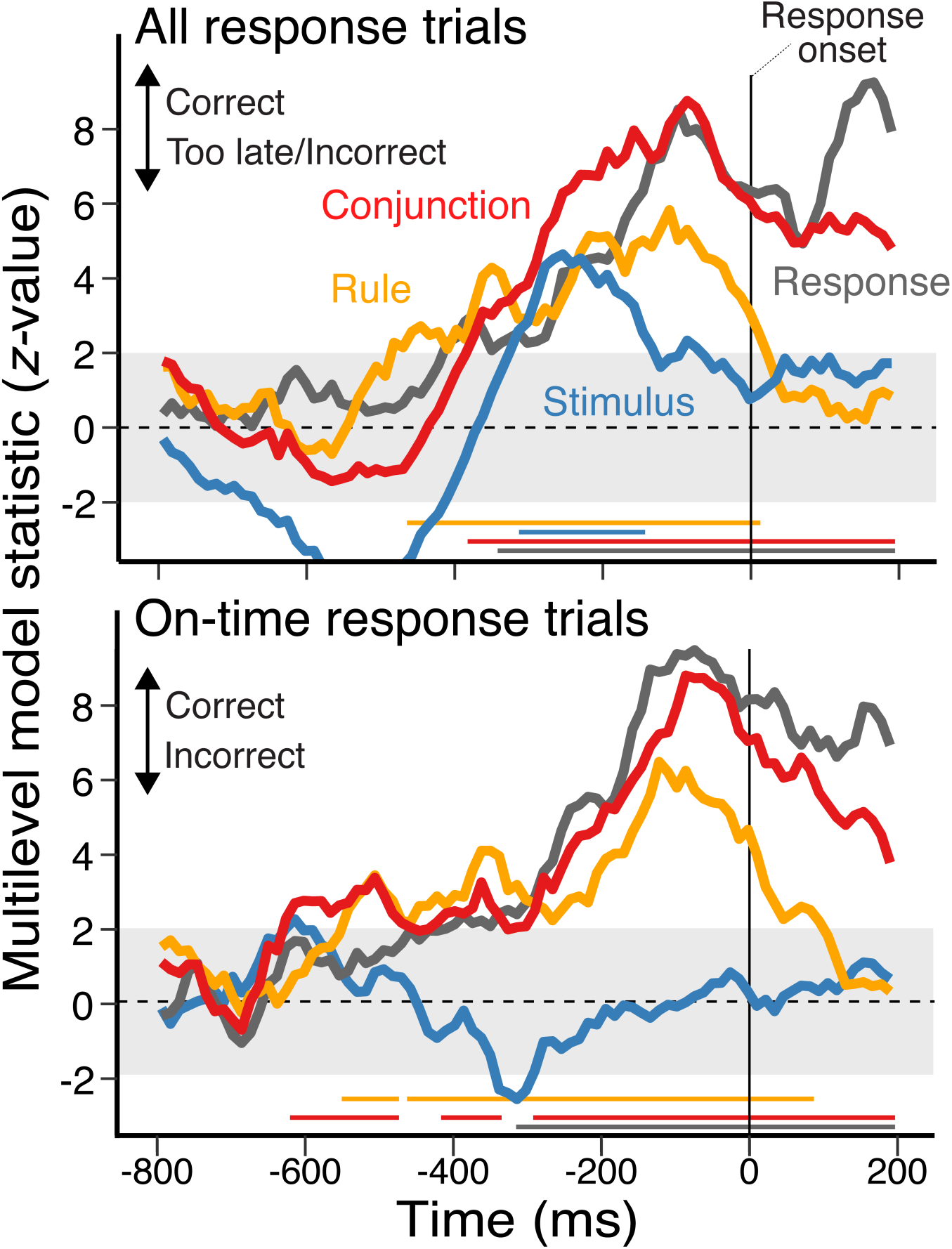
Time-course of impact of decoding of task representations on successful response selection. The time-course of the effect of decoded task representation strength on response selection performance. The *z*-values were obtained from multilevel, logistic regression models predicting the variability in trial-to-trial accurate and on-time responses from the strength of task representations at each moment. The signals are aligned to the onset of trial-to-trial responses. The top panel shows the result using all responses except complete omissions and premature responses, whereas the bottom panel shows the result with responses made within the deadline. Positive *z*-values indicate more successful responses as the strength of decoded representations increases. The colored straight lines at the bottom denote the significant time points using a nonparametric permutation test (cluster-forming threshold, *p* < .01, cluster-significance threshold, *p* < .01, two-tailed).

### Task Representational Dimensionality

How does the representational dimensionality of a control representation change over the course of neural trajectories? How does this relate to successful action selection? And, what are the functional links to the quality of the component task representations (i.e., stimulus, rule, and response) and the conjunctions (Figure 2C)? To this end, we sought to measure the time-dependent changes in the representational dimensionality (i.e., shattering dimension) of task representations. To estimate the dimensionality of the neural responses, we adapted the binary classification method used in physiological non-human primate recording data (Rigotti et al., 2013) for substantially noisier human scalp EEG data by adding constraints on selecting successful separations. This exclusive cutoff method is detailed in the *Dimensionality Analysis via Binary Classification Method*.

The results of the simulation verified that the binary classification method with exclusive cutoff could be applied to noisy data where the separability of patterns is generally low to study the geometry of neural responses (Figure S8). Further, we tested how the selectivity of patterns that are tuned for basis features of the current task in a pure-selective vs. mixed-selective (i.e., conjunctive) manner contributes to the representational dimensionality. Note that, to focus on the geometric properties provided by the integration of information irrespective of the scale of the neural activity (e.g., at the level of single neurons or manifolds), we sampled these patterns directly from RSA models (Figure 1C) to generate patterns that differ in the degree of integration of task-relevant information. We confirmed that in patterns of neural activity that is exclusively tuned to one specific instance of the action constellation like an event-file (i.e., excluding diverse mixing of task features), as assumed in the conjunction RSA model (Figure 2C), the dimension of neural patterns expanded substantially (Figure S9; see *Dimensionality Analysis via Simulation* in the Supplementary Information for detail).

We applied our validated (see supplement) exclusive cutoff pairwise classification method in a time-resolved manner in order to characterize the dynamic changes of representational dimensionality while context-dependent action selection unfolds. Consistent with our hypothesis that the benefits of high dimensional representational geometry on controlled actions by enhancing separability, we found that the dimensionality transiently peaked before response execution and then collapsed rapidly (Figure 6). Consistent with the previous works in non-human primates, we found that the dimensionality is overall significantly higher when participants made correct and on-time responses (Figure 6; *t*(1,41) = 9.09, *beta* = .48, 95% CI [.38 .59] for stimulus-aligned and *t*(1,41) = 7.39, *beta* = .42, 95% CI [.30 .53] for response-aligned). Importantly, the difference appeared even before the trigger for response-deadline was presented, *t*(1,41) = 5.43, *beta* = .37, 95% CI [.24 .51], and persisted until the end of deadline, *t*(1,41) = 6.14, *beta* = .45, 95% CI [.31 .60]. Further, the same effect was observed in the short SOA condition where the number of correct and too late or incorrect responses were matched in principle, *t*(1,41) = 3.59, *beta* = .26, 95% CI [.12 .41] for response-aligned. Both the temporal trajectories and the condition difference of dimensionality were preserved for higher cutoff threshold values (Figure S10BD). Thus, consistent with our hypothesis, representational dimensionality changes dynamically over the course of action selection and its development strongly correlates with more successful implementation of context-dependent actions.

**Figure 6.**
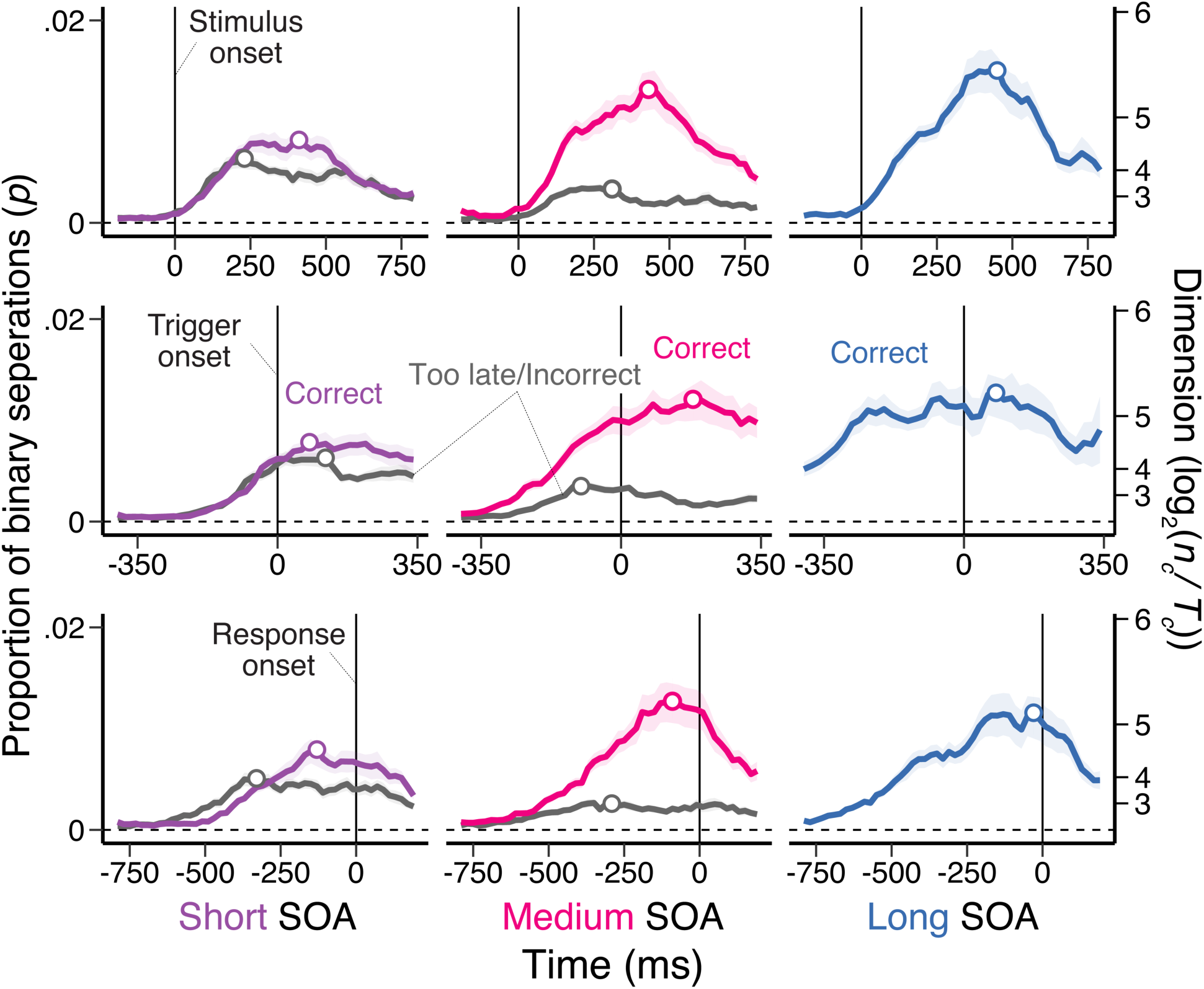
Time-course of task representational dimensionality. Average, dimension score during action selection. The dimensionality is estimated as the proportion of implementable binary separation, *n_c_/T_c_*, out of all possible arbitrary binary pairs. The left, middle and right column corresponds to the short, medium, and long SOA conditions respectively. The top, middle and bottom row correspond to the results using signals aligned to the onset of stimulus, trigger or response in each trial. The two lines in each panel show the average scores in trials with correct vs. too late or incorrect responses. Shaded regions specify within-subject standard errors.

How does the representational dimensionality relate to the quality of action representations? To investigate this, we summarized the result of a binary classification analysis with exclusive cutoff (*p* = .51) separately for decile bins of RSA scores of each task factor across trials. Based on the results of the simulation, we expect a positive relationship between the strength of signal and the representational dimensionality (*n_c_/T_c_*) where the steepness of functions relating to these two measures depends on the tuning profile and the noise level of underlying neural activities (Figure 6). Consistently, we found that higher RSA scores of the rule, stimulus, response, and conjunction overall expand the representational dimensionality (Figure 7). The degree of enhancement was strongest for conjunctive representations, supporting that RSA-derived conjunctions indeed reflect the mixture of task-critical information. The effect of conjunctions on the dimensionality was preserved for higher cutoff threshold values (Figure S10AD). Taken together, these results confirm that higher dimensional representational geometry that accompanies encoding of event-file like conjunctive representations enhance separability for linear readouts, which strongly predict the successful implementation of goal-contingent action selection.

**Figure 7.**
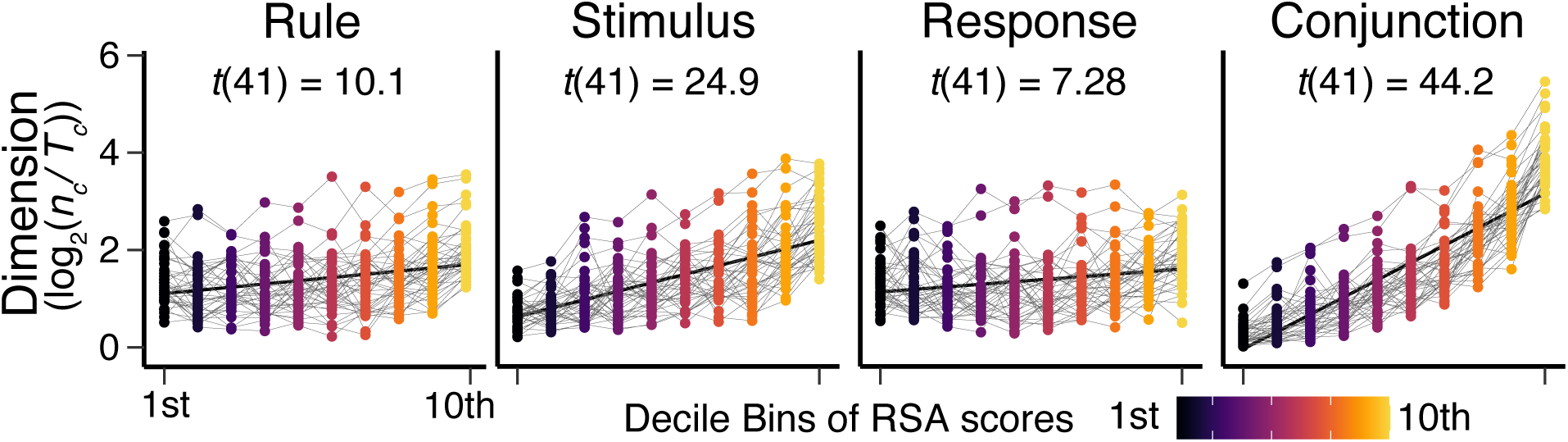
Relationship between the dimensionality and selectivity of task representations. Changes in the representational dimensionality as a function of RSA scores of each of the basis set task features (rule, stimulus, and response) and their conjunction. The dimension scores (*n_c_/T_c_*) are calculated separately in decile bins of RSA scores of each feature within subjects, which correspond to individual points in each panel. The *t*-value is the statistic of a linear regression model fitted to the results.

### Task Representational Dynamics

By focusing on the geometry of neural responses in a time-resolved manner, we found that encoding of conjunctive control representations recruit a transient high-dimensional geometry, which peaks prior to response execution, that enhances separability of neural patterns. Yet, the utility of such heightened separability partially depends on the underlying dynamics. This is because to implement optimal readouts of information, the neural trajectories must be either predictable or stabilized within the projected task relevant subspaces. To assess the stability of underlying neural trajectories, we combined the cross-temporal generalization analysis (Dehaene and King, 2016) with single-trial RSA.

We first performed an exploratory analysis using our previously published data using a similar task without the response deadline, Kikumoto and Mayr, 2020. The result showed that a similar increase of the representational dimensionality that peaked slightly before response execution, which scaled with the strength of conjunctive representations (Figure 6 bottom and Figure S11 top). This result indicates that a response deadline itself is not a necessary condition to induce expansion of representational dimensionality. In addition, we found enhanced separability was established when the conjunctive subspaces became temporarily stable which are indicated by the decision hyperplanes becoming more generalizable to other time points (Figure S11 bottom). In particular, early forward generalization of the decision hyperplane (i.e., upper-left portion of the decoding matrix) preceding the onset of responses suggests that there is an attractor-like stabilization of neural trajectories that will retain readouts of neural states to be generalized for future states.

How the convergence of neural dynamics influences successful action selection is difficult to assess without any constraints on encoding of neural trajectories for task computations. For example, slow responses may reflect failures of readout of task representations or strategically withholding for the stabilization of dynamics. In addition, the passage of time is often confounded with temporal stabilization of dynamics as in the cases in typical delayed maintenance tasks. The current study’s use of the response-deadline procedure circumvents these issues by forcing participants to use the product of dynamic computations at premature or developed states.

By assessing temporal generalizability of linear hyperplanes that separate conjunctions defined by the combination of task basis features, we found that neural trajectories in the conjunctive subspace became relatively stable toward the moment that responses are executed (Figure 8). Importantly, gradual expansion of representational dimensionality peaked while the underlying dynamics were stable when accurate and on-time responses were achieved. In contrast, such temporally stable dynamics emerged at much later time windows when responses were incorrect or too late (see Figure S12 and Figure S13 for the results for other task factors).

**Figure 8.**
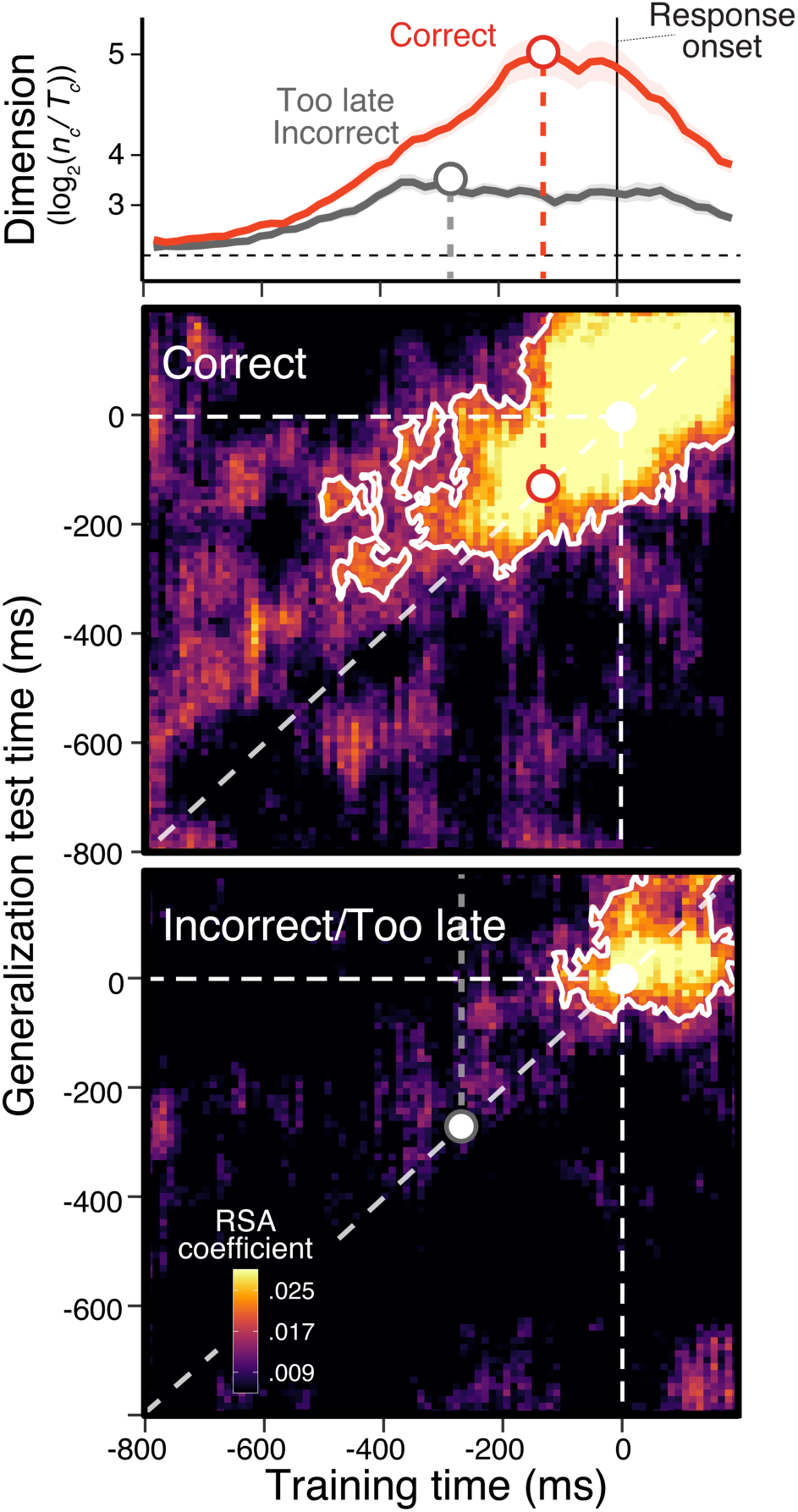
Representational dynamics in the subspace of conjunctive control representations. Temporal generalization of linear hyperplanes that separate conjunctions of task basis features summarized as a decoding matrix. The X-axis denotes time points where signals were sampled to train decoders then submitted to single-trial RSA. The Y-axis denotes time points where the test sets were derived. Thus, the diagonal components of the matrix correspond to trajectories of time-resolved decoding analyses, as shown in Figure. 4. The off-diagonal components show the results of temporal generalization analysis. The regions surrounded by a white contour denote the significant clusters using a nonparametric permutation test (cluster-forming threshold, *p* < .01, cluster-significance threshold, *p* < .01, two-tailed). The top panel shows changes in the representational dimensionality relative to the onset of responses (Figure 7 bottom row) merging short and medium SOA conditions. The middle panel plots temporal generalization for correct responses, and the bottom panel plots temporal generalization for incorrect or too late responses. Shaded regions of the top panel specify within-subject standard errors.

To test the importance of early development of stable dynamics, we predicted single-trial response successes by forward generalization of hyperplanes of all task-relevant features while controlling for predictability of dynamic portion of variability in encoded task representations. Critically, we found that more temporally stable dynamics in subspaces encoding conjunctive representations, over and above the dynamic coding of all task factors and the conjunction, uniquely predict higher probability of selecting responses accurately within the deadline, *t*(1,41) = .21, *beta* = .01, 95% CI [-.03 .04] for rule, *t*(1,41) = 2.65, *beta* = .05, 95% CI [.02 .07] for stimulus, *t*(1,41) = .71, *beta* = .14, 95% CI [-.02 .04] for response, and *t*(1,41) = 2.60, *beta* = .05, 95% CI [.01 .09] for conjunction, using time window from -400 ms to 0 ms relative to the onset of responses (see Supplementary table 4 and Supplementary table 5 for the statistics of all other predictors in the model and the results using both forward and backward generalizations with different sets of time windows). Note that the dynamic component of stimulus encoding did not predict response failures (Figure 5, and Supplementary table 5), thus the effect is likely driven by shifting early dynamic encoding of stimulus further past because of delayed responses (Figure S13). The unique contribution of conjunctive representations were detected when the signals are aligned to the onset of stimuli, *t*(1,41) = 2.15, *beta* = .03, 95% CI [.01 .05] for rule, *t*(1,41) = 1.23, *beta* = .02, 95% CI [-.01 .04] for stimulus, *t*(1,41) = 3.99, *beta* = .06, 95% CI [.03 .08] for response, and *t*(1,41) = 4.18, *beta* = .07, 95% CI [.04 .10] for conjunction, using time window from 0 ms to 400 ms relative to the onset of stimuli.

Taken together, these results suggest that effective controlled actions involve the recruitment of temporally stable conjunctive subspaces. The earlier establishment of these stable conjunctive representations is linked to more efficient controlled behavior, in addition to the benefits derived from their dynamic computations. The stability of these dynamics is attained simultaneously with the transient expansion of the representational dimensionality of neural trajectories, which promotes more reliable readouts while facilitating the separation of critical information for the task at hand.

## Discussion

Flexible, goal-directed behavior depends on reliably integrating specific action pathways with abstract knowledge about goals and contexts. At the heart of this integrative capacity are control representations that select, organize, and maintain task-critical information using neural codes suited to the computational demands of behavior. Control representations must be context-sensitive, separating highly correlated inputs into distinct neural states for efficient readout. Yet, they must also be robust in the face of the inherent temporal variability of noisy, biological computation, allowing for stable readout for flexible behavior. In this study, we investigated how control representations are expressed within the complex, time-varying geometry of neural state space. In particular, we tested the hypothesis that temporally stable conjunctive spaces expressed within a high-dimensional geometry influenced controlled action selection, thus enabling both separability and stability required for cognitive control problems. To this end, we characterized the temporal development of representational dimensionality and subspaces for individual task features relative to behavioral events, adapted the classification-based dimensionality estimation for the noisy neural data in humans, and explicitly related the dimensionality to encoding of conjunctive representations.

First, we found that the relevant contexts or action rules are integrated with other key task information as conjunctive control representations, replicating prior work (Kikumoto & Mayr, 2020). Extending our previous findings, our procedure with variable SOA intervals and a response deadline demonstrated that the encoding of integrated task features was crucial for the efficient execution of controlled actions independent of the decision to make a response. Moreover, we found that reaching a neural state that allows for the temporally stable readout of conjunctions precedes successful responding, regardless of the enforced variability in the timing of response execution. The encoding of conjunctive and response representations was weak or absent when responses were delayed or incorrect, possibly indicating impaired readout of responses (Figure S6).

Second, we found that neural representational (i.e., embedding) dimensionality transiently changed within each episode of action selection. Specifically, during episodes that produce successful, controlled actions, dimensionality rapidly expanded and peaked just prior to response execution, leading to a sudden drop. When this expansion had not yet happened, participants failed to make correct or on-time responses. Furthermore, this higher-dimensional geometry particularly enhanced the conjunctive control representations. These findings align with our simulation results and are in line with theoretical and empirical evidence suggesting that neural mechanisms involved in integrating task-critical information, even by encoding exclusive conjunctions alone, expand the embedding dimension of the representation. During this transient period of dimensionality expansion, the separability of the neural patterns encoding task states is increased, making task-critical features accessible to a linear readout.

Finally, we found that the complex neural trajectories encoding control representations contained both dynamic and temporally stable components. In particular, we found evidence of conjunctive subspaces that exhibit temporal stability in the period just prior to response execution, despite the ongoing dynamics of neural trajectories in the entire state space. The timing of accessing the stable conjunctive state explained unique variance in achieving successful response selection beyond that explained by the strength of the relevant task variables, including conjunctions, decoded from the dynamic trajectories alone. The contributions of both dynamic and stable components were observed in conjunctive subspaces uniquely, suggesting efficient dynamics for task-tailored solutions reflecting required goal states (Stroud, et al. 2023).

Thus, our overall results support the hypothesis that encoding a stable conjunctive subspace within a high-dimensional representational geometry provides the enhanced separability and temporal stability that are ideal for solving cognitive control problems. Several recent theories and their associated computational models of cognitive control (Cole et al., 2013; Frings et al., 2020; Hommel et al., 2001; Ito et al., 2022; Logan, 1989; Schumacher & Hazeltine, 2016) directly emphasized that the integration of task-relevant features is an important computational step toward flexible outputs, which contrasts with the more traditional view of cognitive control recruiting a hierarchical, multi-step process (Miller & Cohen, 2001; O’Reilly & Frank, 2012; Koechlin, Ody & Kouneiher, 2003). Our findings impose additional empirical constraints on these theories, as they must account for an integrative subspace that facilitates the separation of task-critical information in a temporally stable manner, in concert with a dimensionality expansion. Below, we further elaborate on why such features of control representations may benefit the system and the implications of these observations for theory and future work.

Many cognitive control theories assume integration or association of goals with other task-relevant features at some level, but not all emphasize these as the control representation needed to solve context-dependent behavioral problems. For example, the widely influential Cohen, Dunbar & McClelland (1990) guided activation model proposed a solution to the Stroop task via lateral inhibition without units that encode separate conjunctions between task demands and other features. This leads to the view that control is achieved by goal representations maintained in a factorized, low-dimensional (i.e., compositional) format, which biases neural activity in action selection stages. Our results indicate that the encoding of conjunctions is also critical for implementing context-sensitivity, by directly associating goal contexts into stimulus and response bindings (Hommel 2011; Frings et al., 2020; Schumacher & Hazeltine, 2016; Verguts & Notenaert, 2009). They further indicate that encoding stable conjunctions can be understood as a mechanism that recruits additional embedding dimensions to bind goals, contexts, and action rules with other compositional task features.

In this sense, conjunctions may serve the computational function of enabling linear separability of the low dimensional inputs, thereby guiding cognitive control. Indeed, the embedding dimensionality of neural activity adapts to the complexity of the computational demands of the task, often enhancing the separability of compositional input features (Bernardi et al., 2018; Ehrlich & Murray, 2022; Fu et al., 2022; Gao et al., 2017; Ito & Murray, 2023; Jazayeri & Ostojic, 2021b; Park et al., 2023). Similarly, to afford linear separability of low dimensional inputs, recurrent neural networks with chaotic dynamics also show temporally dimensional expansion prior to predictable readout (Farrell et al., 2022). In cognitive control tasks requiring context-dependent mappings, conjunctions play a crucial computational role in disambiguating similar neural states induced by low-dimensional inputs, by projecting them onto a higher-dimensional geometry. Supporting this interpretation, we observed a robust reliance on conjunctions for successful context-dependent actions (Figure 5) and discovered a significant correlation between the strength of conjunctions and the dimensionality of neural activity (Figure 7; Figure S10). Thus, this viewpoint explains how encoding of conjunctive state itself enhances the separability of control representations in the service of context-sensitivity by inducing higher embedding dimensions. This connection between the dimensionality to a theoretically important control representation has not been established explicitly in humans.

Further, we observed that the transient expansion of the embedding dimension increased beyond the dimensionality strictly required to solve the task, supporting more diverse linear separations of task variables than needed. Such an expansion of the embedding dimension, which consistently occurred just prior to response execution, should be beneficial for creating additional subspaces by pushing neural patterns closer to orthogonality (Rigotti et al., 2013). However, the observed transient, higher-dimensional geometry is not solely explained by the encoding of conjunctions, as suggested by the mismatch between the time courses of decoded conjunctive representations and dimensional expansion (Figure 4 and Figure S11), and the lower average level of dimensionality even in trials with strong conjunctive representations (Figure 7). One possible advantage of a larger embedding dimension at the moment of transition to response execution, is to accommodate both the response and the conjunctive subspaces, thereby achieving better dissociation of neural trajectories. Similarly, several recent studies in non-human animals have observed the dynamic use of orthogonal subspaces to separate different task-critical information, potentially reducing interference of behaviorally relevant variables (Bagur et al., 2022; Elsayed et al., 2016; Fine et al., 2022; Libby & Buschman, 2021; Wan et al., 2022; Weber et al., 2022). An important area for future research is to clarify the neural mechanisms that generate transient dimension expansion and their relationship to stable subspaces of task representations.

Our results also emphasize that control mechanisms implement a temporally stable yet integrative subspace within dynamic neural trajectories. Such solutions with stable subspaces not only enable reliable readout of information but also provide a few advantages in solving cognitive control problems. In neural networks, optimizing for context-sensitivity often induces highly dynamic neural trajectories as solutions (Buonomano & Maass, 2009; Farrell et al., 2022). A computational consequence of this time-varying activity is that the readout must learn distinct weights for each moment. In contrast, encoding a stable conjunctive subspace within high dimensional geometry allows for a fixed, common set of weights that can produce time-invariant readout, while affording the context-sensitivity required by cognitive control. Such stable subspaces provide flexible and reliable readout over variable temporal delays until the completion of the tasks (Cueva et al., 2020; Murray et al., 2017). This temporal invariance provides the ability to decode task-critical information (e.g., to select a response) at arbitrary timings without explicitly encoding time, and it also offers robustness to the time-varying nature of noisy computation afforded by biological circuits. Though we note that the periods of stability we observe here were brief, arising in the moments prior to the response. So, whether they would bridge longer variable delays is a matter for future research. This caveat notwithstanding, however, a stable state within conjunctive subspaces also promotes learning a single set of readout weights despite the dynamic nature of the overall neural trajectory. This significantly reduces the complexity of learning downstream readouts, facilitating more efficient learning of the task.

Our results add to the converging evidence that costs on behavior directly emerge from the dynamics and geometry of task representations. Various psychological and neural factors such as the objectives and states of learning, task demands, and interactions of multiple cognitive systems could influence these properties.

For instance, gaining expertise in a fixed task environment has been theorized to reflect more efficient encoding or retrieval of instances or task-specific memory that minimally specifies required goal states (Logan, 1988). Given that stabilization and expansion of dimensions is not achieved instantaneously, how efficiently the system reaches highly separable geometry and a stable, conjunctive state may be directly optimized through experience. The frequently observed co-existence of initially dynamic and later stabilizing subspaces of task representations may reflect the efficient transformation of task information in a goal-contingent manner (Myers, 2022; van Ede et al., 2019; van Ede & Nobre, 2022). In that case, the benefits and costs in response selection such as repetition priming (Benini et al., 2022; Hommel et al., 2011; Mayr & Bryck, 2005), pre-cueing preparation effects (Altman, 2004), or multi-tasking costs (Marti et al., 2015), could also depend on how the dynamics and geometry unfold in the relevant control episodes (Kikumoto & Mayr, 2020; Rangel et al., 2023).

A limitation of the current study is that our analysis of EEG data cannot inform our understanding of the scale and locus of the neural computations underlying conjunctive subspaces and their readout. This ambiguity arises from the difficulty in distinguishing between connectivity and activity of neural assemblies at the level of EEG. The signals we observe may reflect local computations within specific brain regions and networks (Asaad et al., 2000; Lapate et al., 2021; Rigotti et al., 2013; Stokes et al., 2013; Weber et al., 2022) or macroscale brain states resulting from distributed interactions among multiple neural populations and networks (Buschman et al., 2012; Greene et al., 2023; Gu et al., 2015; MacDowell et al., 2022; van den Brink et al., 2022). Thus, future research should aim to distinguish these alternatives, perhaps through complementary methods in humans and animals that combine neural dynamics with spatially resolved data (Badre et al., 2015). However, we note that our representational similarity analysis using a GLM with competing models makes it is unlikely that conjunctions arise from stimulus, rule, and response information co-occurring in separate pools of domain-specific electrodes (Kikumoto & Mayr, 2020).

Another limitation is how generalizable our findings are beyond the current task, where context-sensitivity is explicitly required to solve the task. Our claim is that enhanced, stable separability in task representations – which is partially provided by the encoding of stable conjunctive subspaces – is useful for performing any task requiring context sensitivity. This kind of context specificity (i.e., mapping the identical inputs to different responses depending on the contexts) is a feature of many, if not most, standard cognitive control tasks such as the Stroop task, task-switching tasks, the Wisconsin Card Sorting task, S-R compatibility tasks, the Flanker task, the MSIT, etc. However, it is also possible that the geometric and dynamical complexity of task representations adapts flexibly to task demands in a brain region-specific manner (Mack et al., 2020; Bhandari et al., 2024). The generalizability of our findings, such as stable conjunctive subspaces and transient dimensionality expansion, should be directly empirically tested in future studies.

In conclusion, we characterized the geometric properties and dynamics of control representations that facilitate separability and stability for reliable and flexible action selection. Specifically, when people successfully perform context-dependent action selection, representational dimensionality expands, coinciding with temporal stability in subspaces defined by conjunctions of key task features. Early entry into this stable state and enhanced separability are crucial for efficient behavior. These findings prompt a new mechanistic focus on cognitive control, geared toward understanding the conditions and processes that implement and bring about these representational states. And they underscore the potential value of adopting both a geometric and a dynamic state-space perspective to gain deeper insights into the control representations that govern our adaptable behavior.

## Methods

### Participants

Forty-two participants (27 female; mean age: 22 years) were recruited and gave informed consent. This followed procedures approved by the Human Subjects Committee at the RIKEN. The sex and gender of participants were determined based on self-report. No sex and gender analysis was carried out because our hypotheses were not dependent on it. No statistical method was used to predetermine sample size. All participants had normal or corrected-to-normal vision and had no history of neurological or psychiatric disorders. After preprocessing the EEG data, one participant was removed and was not analyzed further due to excessive amounts of artifacts (i.e., more than 25% of trials; see *EEG recordings and preprocessing* for details).

### Behavioral Procedure

Throughout all phases of the experiment (see below), participants performed a rule-based action selection task (Mayr and Bryck 2005; Figure 2A), which required participants to behave according to 12 unique stimulus-response (S-R) mappings (Figure 2B). The twelve S-R mappings were defined by the combination of three spatial transformation rules (vertical, horizontal and diagonal) over four stimuli (a dot at top-left, top-right, bottom-left, and bottom-right of the white frame). The rules described how to select one of four responses that were arranged in 2 x 2 square matrix (4, 5, 1, and 2 on the number pad) based on the stimulus. Specifically, the diagonal rule required the participant to choose the response that was diagonally adjacent to the stimulus location. For example, if a dot was presented in the left-bottom of the frame and the vertical rule was cued, this specified the left-top response as correct (Figure 2A). Accordingly, the vertical rule specified the vertically adjacent response, and the diagonal rule specified the diagonally adjacent response (Figure 2B).

Thus, for each trial, participants selected one of the four responses based on a random cue for one of the three action rules and one of the four stimuli, yielding one of 12 unique combinations of rule, stimulus, and response. One experimental block lasted 18 seconds, during which participants were instructed to complete as many trials as possible. Trials that were initiated within the 18 second block duration but extended beyond it were allowed to finish. There were 25, 35, and 185 experimental blocks for each task phase. Before participants started the no-deadline task, they practiced the same baseline task (i.e., without deadlines) for 15 blocks.

Participants performed this rule-based selection task in three task phases in order: the no-deadline phase, the variable stimulus-onset asynchrony (SOA) phase, and the sampled SOA phase. In the no-deadline phase, after 15 practice blocks, participants performed the rule-based action selection task and were only instructed to respond as accurately and quickly as possible, balancing both equally. The no-deadline phase provided reference RTs, which were used to adjust the timing of SOA intervals for subsequent phases as described in the Estimation of SOA functions (Figure S1A).

In the variable SOA task, participants were presented an imperative auditory stimulus as the trigger signal, at a point following the stimulus and rule presentation (Figure 2A). They were required to make their response within 350ms of hearing the trigger signal. The trigger signal (a pure tone in 500 Hz and 250 ms duration) was presented after one of 12 unique, subject-specific SOA intervals, where SOA was defined as the time interval between the stimulus-rule presentation and the trigger sound. One of these 12 unique SOAs was sampled randomly for each trial. The determination of timing of the SOA is described below. Importantly, here participants were instructed and regularly encouraged to make prompt responses within the deadline following the trigger signal even if they are not ready to respond.

In the sampled SOA phase, we measured EEG while participants performed the same task but with three SOA intervals that were estimated from SOA functions, defined from the variable SOA phase, to produce response errors (including responses that are incorrect and/or too late) at 50%, 70% and 90% of trials. The SOA intervals of each error probability condition are denoted as short, medium and long SOA respectively. One of these 3 unique SOAs was sampled at random for each trial.

### Estimation of SOA functions

To sample responses at fixed error probabilities during the EEG session, we estimated individual SOA psychophysical functions by systematically modulating SOA intervals before the deadline (“trigger”) signal (Figure 2A). We first obtained the distribution of RTs of correct responses in the no-deadline phase excluding trials where RTs were slower than 5 SD. The RT distribution was modeled by the ex-Gaussian distribution using the exgauss toolbox in Matlab 2019B (Zandbelt 2014; Figure S1B). Then, in the variable SOA phase, participants performed the same task with 12 SOA intervals ranging from -450 ms to 200 ms (-450, -400, -350, -300, - 250, -200, -100,-50, 0, 100, 200 ms) referenced to their ex-Gaussian μ parameters. Negative SOA intervals were dropped during estimation. As a result, we obtained individuals’ probabilities of producing accurate and on-time responses as a function of SOAs averaged over trials (Figure 3A).

Following (Öztekin & McElree, 2010), we estimated the changes in the rate of successful response selection in variable SOAs with an exponential function approaching to a limit:

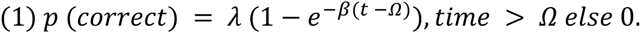

In the equation above, t is the predicted p(correct) or the probability of correct responses at SOA interval t; *λ* is the asymptotic accuracy level reflecting the overall probability of successful responses; *β* is the rate parameter, which indexes the speed at which accuracy grows to asymptote; and *Ω* is the intercept reflecting the discrete point when accuracy departs from 0. The parameters were estimated by nonlinear least square solver via a lsqcurvefit function in Matlab, which performs nonlinear curve fitting using a least square solution with the lower bound of 0, 0, and the minimum value of tested SOA intervals; and the upper bound of 1, 1, and the maximum value of tested SOA intervals. Note that the sole purpose of the estimation of SOA functions is to induce sufficient number of response errors consistently across participants, thus we do not intend to propose the SOA model reflects underlying cognitive processes. Using this approach, we were able to predict the probabilities of correct responses as a function of continuous SOAs (Figure 3B and Figure 3C) to produce expected distributions of errors (Figure 3D).

### EEG recordings and preprocessing

The continuous EEG was recorded using a Brain Products actiCHamp recording system (Brain Products GmbH). Recordings were obtained from a broad set of scalp sites (Fp1, Fp2, F7, F3, F4, F8, FC5, FC1, FC2, FC6, T7, C3, C4, T8, CP5, CP1, CP2, CP6, P7, P3, P4, P8, PO9, PO10, O1, O2, Fz, FCz, Cz, Pz, and Oz), from the left and right mastoids, and from electrodes lateral to the external canthi and below the left eye. The electrodes (TP9 and TP10) were placed on the mastoids beneath the cap as reference sites. Using additional passive electrodes, horizontal electrooculogram (EOG) was recorded from laterally placed electrodes to detect horizontal eye movements with the separate ground electrode placed on the left side of forehead. We verified that eyeblinks were visible and recorded at frontal electrodes such as Fp1 and Fp2. The EEG was filtered online with .1 Hz high-pass and 1000 Hz low-pass filters and digitized at 1000 Hz. The left-mastoid was used as reference for all recording sites, and data were re-referenced off-line to the average of left and right mastoids. The EEG data were obtained during the sampled SOA task after participants-specific SOA intervals were estimated.

The scalp EEG and EOG were amplified with an SA Instrumentation amplifier with a bandpass of .01–45 Hz, and signals were downsampled at 250 Hz using EEGLab (Delorme & Makeig, 2004). EEG data was first segmented into 22 second intervals to include all trials within a block and to remove edge artifacts in later analysis steps. After time-frequency decomposition was performed, these epochs were further segmented into smaller epochs for each trial. For stimulus-aligned epochs we used the time interval of -200 ms to 800 ms relative to the onset of stimuli, which encompasses all SOAs of all participants (Figure 3A). For response-aligned epochs we used the time interval of -800 ms to 200 ms relative to the onset of responses. For epochs aligned to the response deadline signal (i.e., SOA-aligned), we re-epoched intervals from the stimulus aligned epochs using the time interval of -400 ms to 350 ms relative to the onset of a signal reflecting participants specific SOA intervals. The trial-to-trial epochs that included blinks (>250 μv, window size = 200 ms, window step = 50 ms), large eye movements (>1°, window size = 200 ms, window step = 10 ms), blocking of signals (range = -.01 to .01 μv, window size = 200 ms) were excluded from subsequent analyses. Any epochs that showed EEG activities larger than 5 s.d were excluded.

### Time-Frequency Analysis

Temporal-spectral profiles of single-trial EEG data were computed via complex wavelet analysis (M. X. Cohen, 2014) by applying time-frequency analysis to preprocessed EEG data epoched for the entire block (>22 seconds to exclude the edge artifacts). The power spectrum was convolved with a series of complex Morlet wavelets ((2) *e^i2πft^ e*^−(*t*/*σ*)2^/2), where *t* is time, *f* is frequency increased from 2 to 40 Hz in 35 logarithmically spaced steps, and σ defines the width of each frequency band, set according to *n*/2π*f*, where *n* increased from 3 to 10. We used logarithmic scaling to keep the width across frequency band approximately equal, and the incremental number of wavelet cycles was used to balance temporal and frequency precision as a function of frequency of the wavelet. After convolution was performed in the frequency-domain, we took an inverse of the Fourier transform, resulting in complex signals in the time-domain. A frequency band-specific estimate at each time point was defined as the squared magnitude of the convolved signal for instantaneous power.

### Representational Similarity Analysis

We performed a time-resolved multivariate pattern classification analysis to decode action-relevant information following our previously reported method based on the representational similarity analysis with a few modifications (Kikumoto & Mayr, 2020; Kriegeskorte et al., 2008). As the first step, at every sample in trial-to-trial epochs, separate linear decoders were trained to classify twelve possible action constellations or each of S-R mapping (Figure 2B). We used a penalized linear discriminant analysis using the caret package in R version 4.3.1 (Hastie et al., 1995; Kuhn, 2013) to solve this multiclass classification problem. This step produced a graded profile of classification probabilities for each action constellation over time, reflecting the similarity of underlying multivariate neural patterns between actions. For example, in trials where the horizontal rule and the left-top stimulus were presented, the goal of decoders is to assign a higher probability for that specific S-R condition over others (Figure 2C). The same target label is used for all time points in the same trials.

Decoders were trained with the instantaneous power of rhythmic EEG activity, which was averaged within the predefined ranges of frequency values (2-3 Hz for the delta-band, 4-7 Hz for the theta-band, 8-12 Hz for the alpha-band, 13-30 Hz for the beta-band, 31-35 Hz for the gamma-band), corresponding to 155 features (5 frequency-bands X 31 electrodes) to learn. Prior to the training of decoders, trials where responses occurred before the deadline period (i.e., before the trigger signal) and trials where responses were completely omitted were excluded. Further, all trials in the first block (of the sampled SOA task phase) were excluded.

Within individuals and frequency-bands, the data points were z-transformed across electrodes to remove effects that scaled all electrodes uniformly. We used a k-fold repeated, cross-validation procedure to evaluate the decoding results (Mosteller & Tukey, 1968) by randomly partitioning single-trial EEG data into five independent folds with an equal number of observations of each action constellation. After all folds served as the test sets, each cross-validation cycle was repeated ten times, in which each step generated a new set of randomized folds. The number of observations in each target class (e.g., action constellation) was equated by randomly dropping excess trials for certain conditions. In addition, the effect of subsampling the number of trials for correct, incorrect, and too late responses was assessed in separate analyses (Figure S3). Resulting classification probabilities were averaged across all cross-validated results with the best-tuned hyperparameter to regularize the coefficients for the linear discriminant analysis.

We next performed representational similarity analysis (RSA) on the graded classification probabilities to assess the underlying similarity structure of task variables. Each RSA model matrix uniquely represents a potential, underlying representation (i.e., rules, stimuli, responses, and conjunctions), which makes unique predictions of similarity patterns for different action constellations. To obtain time-resolved estimates of each RSA model, we regressed the vector of logit-transformed classification probabilities onto RSA model vectors for independent decoding results.

Importantly, to estimate the unique variance explained by competing models, we regressed all model vectors simultaneously. The features used for training decoders range in spatial- and frequency-domain, which leaves the possibility of identifying clusters of features that consistently contribute to the decoding of any task features. However, by using competing models, it is unlikely that conjunctions arise from linearly additive effects of co-occurring stimulus, rule, and response information. We also included subject-specific regressors of z-scored, average RTs, and probabilities of on-time and correct responses in each action constellation to reduce potential biases in decoding. These coefficients, expressed in their corresponding t-values, reflect the quality of action representations at the level of single trials, which was later related to variability in behavior and dimensionality measures. We excluded t-values that exceeded 5 SDs from means for each sample point, which excluded less than 1% of the entire samples.

These analysis steps were repeated separately using EEG signals that are stimulus-aligned and response-aligned. The results aligned to the onset of the deadline signal are obtained by sorting stimulus-aligned results to idiosyncratic SOA timings.

### Dimensionality Analysis via Binary Classification Method

To assess the representational dimensionality (i.e., shattering dimension) of action representations, we used a binary (pairwise) classification method (Bernardi et al., 2018; Rigotti et al., 2013). A particular point in the time-series of EEG activities obtained in unique action contexts takes a certain location in trajectories that traverse a multi-dimensional space. The binary classification method evaluates the geometry of representations by counting how many different input conditions, which are defined by the combination of orthogonal action features, could be reliably separated (i.e., shattered) by linear readouts. Specifically, the count of linear separations can be obtained by training a set of linear classifiers that learn to decode all possible arbitrary binary groupings given the input conditions. The number of implementable linear separations is known to scale with the dimensionality of underlying neural activity, which has been closely linked to the nonlinear mixed selectivity like tuning profile of information.

To evaluate the representational dimensionality with EEG data, we performed a binary classification analysis with cross-validation in a time-resolved manner. Rigotti and colleagues (2013) showed that the count of implementable arbitrary binary classifications is robustly related to the dimensionality of the neural responses. Intuitively, distinct neural activity patterns with low amount of noise corresponding to the *c* different input (i.e., task) conditions lead to higher discriminability of patterns, hence, the higher dimensionality of neural state space. The authors identified the point (*c\**) where the fraction of implementable classification (*n_c_/T_c_*) undergoes a phase transition point where the probability of successful binary classifications starts sharply dropping as the sampled input condition *c* increases. The critical value (*c\**) is defined as the maximal value of input conditions (*c*) for which almost all of the binary classifications out of the all possible ways of binary groupings (*T_c_*) could be implemented. This requires an arbitrary cutoff threshold of classification accuracy that is sufficiently above chance level (e.g., *p* = .75-.80 in the original study) and another cutoff threshold for the proportion of *n_c_/T_c_* to determine a sharp dropoff point.

Our goal was to assess the relative differences in dimensionality scores across conditions (e.g., correct vs. incorrect/too-late responses) that scales with the overall separability of neural patterns of EEG that contain a non-trivial level of noise. This requires a lower cutoff threshold of classification accuracy without losing sensitivity to changes in *n_c_/T_c_* driven by expansion or contraction of the dimensionality of neural patterns. To this end, we adapted the original binary classification method introduced in Rigotti et al. (2013) to EEG data by adding constraints on how we select implementable binary classifications (see also the *Dimensionality Analysis via Simulation* in Supplementary information).

Specifically, we trained separate decoders for all possible 4096 (i.e., 2^12^ - 2, excluding two cases where all labels being identical for all *c* = 12 task conditions) sets of arbitrary binary separations defined by 12 unique action constellations or the input conditions (Figure 2B). Resulting decoding accuracies were counted as reliable linear separations when the following two conditions were satisfied: 1) the decoding accuracies exceeded the arbitrary chosen cutoff threshold of decoding accuracy of .51 (Figure 6-7) or higher (Figure S10) where the chance level is .50 and 2) above-threshold decoding accuracies were observed in each of the unique conditions of the input space exclusively (exclusive cutoff method) or in an aggregated manner averaging across all conditions (overall cutoff method). Specifically, the decoding accuracies were first averaged, separately for each action constellation, then a cutoff was applied. This second criterion was applied to increase the sensitivity to detect changes in dimensionality scores driven by the non-linear mixing of information even when the cutoff threshold accuracy is significantly lowered to the chance level (Figure 6). Note that the effect of lowered cutoff threshold values was tested in the simulated data (Figure 6; Figure S8). These results showed showed this approach avoids an inflationary bias on the estimated number of dimensions when using the binary classification method with a low cutoff threshold of decoding accuracy to counteract substantially noisy data (e.g., EEG).

As a result, we obtained the proportion of reliable binary separations out of all possible groupings (i.e., *n_c_/T_c_* = count of implementable binary classifications / (2^12^ - 2)), which approximates the dimensionality of neural responses to the input space (i.e., dimension = log_2_(*n_c_/T_c_*)). When comparing experimental conditions, the aforementioned exclusive cutoff was applied separately in conditions of interest (e.g., SOA or types of responses). The other settings of decoding analyses such as the algorithm of classification, the steps of cross-validation, and the exclusion criteria of trials were identical to the RSA analysis except for that EEG signals being averaged over 20 ms non-overlapping consecutive time windows. The results of stimulus-aligned, response-aligned and go-signal-aligned data were generated in a similar manner as the RSA step.

To test the effect of changing values of the cutoff threshold, we conducted control analyses on both simulated (Figure S8 & S9) and empirical data (Figure S10). We found that increasing cutoff threshold values did not affect differences in dimensionality across critical conditions in empirical data, including their temporal changes, the effect of response accuracy types, and the relationships to decoded task variables (Figure S10). Furthermore, the chosen cutoff threshold replicated the reported main result when applied to the data in Kikumoto & Mayr (2022), where an analogous task without the response deadline was used (Figure S11; see *Dimensionality Analysis via Simulation and Modulation of Cutoff Threshold Values* for detail).

### Temporal-generalization analysis

To assess temporal stability of linear readout or linear hyperplanes learned by decoders, we used the temporal generalization analysis (King & Dehaene, 2016). Firstly, we trained a set of decoders to classify action constellations, following the identical procedure as the time-resolved RSA analysis, using EEG patterns at the specific time point. These time-specific decoders were applied to the held out data obtained at identical (matched) or different (generalized) time intervals. This generates a series of decoding results in a two-dimensional matrix form where the on-diagonal entries correspond to the results in which the time points of the train and test dataset match whereas the off-diagonal entries are the results of generalization of linear hyperplanes over other time. Assessing the pattern of temporal generalization reveals the dynamic nature of underlying neural computation such as the temporal stability of linear readouts of task critical information. The resulting decoding results, corresponding to every pairings of training and testing non-overlapping time-intervals (window size = 12 ms), were submitted to the regression-based RSA procedure to decode rules, stimulus, response and conjunctions simultaneously. The results of stimulus-aligned, response-aligned and trigger-aligned data were generated in a similar manner as in the RSA steps.

### Multilevel Modeling and Permutation Tests

We used multilevel models to further assess the statistical significance of representational dimensionality and decoded representations via RSA and to relate these scores to trial-to-trial variability in behavior (e.g., probability of making on-time responses). We tested statistical significance by either computing time-averaged scores over *a priori* selected time intervals or time-resolved scores combined with non-parametric permutation tests. For models predicting accuracies (i.e., correct responses made within the deadline period), we used a multilevel logistic regression excluding trials with responses that were omitted and/or occurred after the deadline. For models including the dimension scores, only random intercepts are included because the exclusive cutoff method requires aggregation of multiple trials (see *Dimensionality Analysis via Binary Classification Method* for detail). Unless otherwise indicated, subject-specific intercept and slopes were included as the random effects based on trials and participants as independent levels.

For models using time-averaged scores, the intervals were defined as 0 to 500 ms from the onset of stimuli for stimulus-aligned results, -500 to 0 ms from the onset of response for response-aligned results, -400 to 0 ms, and 0 to 350 ms relative to the onset of response-deadline (a trigger signal) for SOA-aligned results. Separate models were constructed using RSA scores averaged over these intervals as dependent variables (Supplementary table 2 and Supplementary table 3) or as predictors (Figure 5 and Figure S7).

For models using time-resolved scores, we performed non-parametric permutation tests on the results of multilevel models (Maris & Oostenveld, 2007). First, we performed a series of regression analyses in the continuous temporal space of interest and identified samples that exceeded threshold for cluster identification (cluster-forming threshold, *p* < .01). For example, for the temporal generalization analysis (Figure 8, Figure S11, S12, and S13), we performed tests in two-dimensional space defined by the training and test time intervals. Then, empirical cluster-level statistics were obtained by taking the sum of *t*-values in each identified cluster with consecutive temporal regions. Finally, nonparametric statistical tests were performed by computing a cluster-level *P*-value (cluster-significance threshold, *p* < .01, two-tailed) from the distributions of cluster-level statistics, which were obtained by Monte Carlo iterations of regression analysis with shuffled condition labels.

## Data Availability

The raw and processed behavioral and EEG data generated in this study have been deposited in the CBS Data Sharing Platform at Human Cognition and Learning under accession code “conjunctions” : https://neurodata.riken.jp/r/Shibata/A%20Transient%20High-dimensional%20Geometry%20Affords%20Stable%20Conjunctive%20Subspaces%20for%20Efficient%20Cognitive%20Control%20(latest)/. The processed data are available at the same site or upon request. The code for behavioral and EEG data analyses are available at the same site or upon request.

## Code Availability

The code used in this study have been deposited in the CBS Data Sharing Platform at Human Cognition and Learning [https://neurodata.riken.jp/r/Shibata/A%20Transient%20High-dimensional%20Geometry%20Affords%20Stable%20Conjunctive%20Subspaces%20for%20Efficient%20Cognitive%20Control%20(latest)/].

## Acknowledgments

We would like to acknowledge the members of the Human Cognition and Learning lab at RIKEN, particularly Takahiro Nishio, Honma Saki, and Sara Matsui. Further, we would like to acknowledge the members of the Badre lab, the laboratory of neural computation and cognition, Learning Memory & Decision lab and the Shenhav lab, particularly Matthew Nassar, Daniel Scott, Alexander Fengler, Harrison Ritz, and Ivan Grahek for helpful comments and discussion. This project was supported by funding from the National Institute of Mental Health (R01 MH125497), the National Institute of Neurological Disorders and Stroke (R21 NS108380), and a Multidisciplinary University Research Initiative award from the Office of Naval Research (N00014-16-1-2832) to DB and from JSPS KAKENHI Grant Number 19H01041, 20H05715, JST Moonshot R&D JPMJMS2013 to KS and the JSPS Overseas Research Fellowships to AK.

## Author Contributions

Project conception: A.K, A.B, and D.B; Methodology: A.K and A.B; Data Collection: A.K and K.S.; Analysis and Interpretation: A.K, A.B, and D.B.; Manuscript writing: A.K, A.B, K.S, and D.B; Funding acquisition: K.S and D.B; Supervision: K.S and D.B.

## Competing Interests

The authors declare no competing interests.

## Supplementary Methods

### Task performance in the no-deadline phase

In the non-deadline phase of the experiment, participants performed the rule-based selection task without the response deadline task. To calculate the reference reaction times (RTs) for adjusting the timing of SOA intervals, response error trials and trials with RTs slower than 5 standard deviations (SD) were excluded from the calculation. The RT distribution was modeled using the ex-Gaussian distribution with three parameters: mean (μ), standard deviation (σ) of the normal distribution, and skewness (τ) of the exponential distribution. The modeling was performed using the exgauss toolbox in Matlab (Zandbelt, 2014). Due to the absence of the cue-stimulus interval, it was expected that the individuals’ mean RT distribution would be slower compared to a previous study (Kikumoto & Mayr, 2020). However, the variability of mean RTs was small (Figure S1A). The subsequent estimation of SOA showed robustness against the variability in the obtained parameters (Figure 3A), as indicated by the individuals’ SOA curves (Figure S2B and S2C). During the sampled SOA phase, the probability of response errors was calibrated to match the expected error rates (Figure 3D).

### Response-aligned RSA trajectories with Subsampling Response Types

The timing of SOA intervals was adjusted based on the probability of correct responses against any other invalid responses. Invalid responses include incorrect responses (commission errors) that occur before or after deadlines and omission of responses (omission errors). The distribution of each response type is summarized in the Supplementary Figure. 2. As expected, both probabilities of too late and incorrect responses as SOA get shorter (see also Table 1).

Because the number of trials of response types that were included in decoding analyses differs substantially, we performed single-trial RSA by subsampling to equate the number of observations of on-time correct, on-time incorrect, and too-late correct/incorrect responses within each fold. After five independent folds with an equal number of observations of these response types served as a test set, each cross-validation cycle was repeated twenty times to obtain coefficients in individual trials included. The resulting strength and temporal trajectories of task variables, relative to stimuli, triggers, and responses, remained largely identical to the original without subsampling, including conjunctive representations (Supplementary Figure.3; see also Supplementary Figure. 6 for further analyses). This pattern of results suggests that worse encoding of rules and conjunctions led to slow responses or incorrect responses, where the cued response information is absent in incorrect trials. Because incorrect and too late responses were led by the similar patterns of development of task representations, in particular for conjunctive representations, we averaged these two conditions as too late/incorrect trials unless mentioned specifically (Figure. 5).

### Stimulus-aligned and Trigger-aligned RSA trajectories

In our previous studies, we employed a similar method for time-resolved decoding of task-related factors via RSA and discovered a gradual unfolding of the cascade of task representations over time. In the current study, we were able to replicate a similar pattern of activity, where the rule and stimulus were followed by the conjunction and then by the response (Figure S4). We observed that conjunctive representations emerged approximately 200 ms after the stimulus onset. Moreover, consistent with the outcomes of multi-level models predicting trial-by-trial response failures, we found that the decoding of rules, responses, and conjunctions was more successful in trials that resulted in correct responses, compared to those that were late and/or incorrect. The trajectories of neural activity also diverged between these two conditions well before the response deadline (Figure S5). These findings not only support previous discoveries indicating that better encoding of conjunctions signifies a higher inclination to execute context-dependent actions (Kikumoto, Samejima, & Mayr, 2021; Kikumoto, Mayr & Badre, 2022) but also suggest that such effects persist even when responses and neural trajectories are time-limited, and the number of successful and failed trials is matched (i.e., in short SOA).

### RSA Trajectories Separating Incorrect and Too Late Trials

Incorrect and too late responses (i.e., those after deadlines) trials were averaged to following the SOA calibration procedure. Yet, it is possible that too late but correct trials and incorrect trials differ in neural trajectories because they reflect different instances of task failure. Thus, we assessed how trajectories differ between these two types of responses after subsampling the number of trials of correct, incorrect, and too-late response trials (see

### Response-aligned RSA trajectories with Subsampling Response Types)

We found that in both too late and incorrect trials, rule, response, and conjunction representations are encoded more weakly compared to correct trials (Supplementary Figure.6). Representations of rules and responses are significantly worse or absent in incorrect trials compared to too-late trials. In contrast, conjunctions were equally worse in these trials up to response execution. These patterns of the results are generally consistent with previous studies where stronger task representations, including conjunctive representations, were predictive of faster response times (Kikumoto & Mayr, 2020; Kikumoto, Mayr & Badre, 2022).

### Predicting Accurate Responses within Deadline Separately in Separate SOAs

The encoding of conjunctions accounted for unique variance in predicting trial-wise responses that were either too late or incorrect (Figure 5, top) and responses that were on-time but incorrect (Figure 5, bottom). To analyze this, we aligned the signals to the onset of trial-to-trial responses, merging subject-specific short and medium SOA intervals, and subjected them to multilevel models. Similarly, we conducted identical analyses by separating short and medium SOA conditions and realigning the signals to the onset of the response deadline trigger. We discovered that the impacts of decoded task variables, including conjunctions, became significant before the response deadline (Figure S7). Notably, the identified clusters of rule and conjunctive representation impacts extended to earlier time samples in the medium SOA condition.

We explicitly tested the difference between pre- and post-trigger period of trajectories coding task variables and its interaction with changes of SOA using multi-level model. In the model, we included the main effect of phase (i.e., pre vs. post-trigger), the linear and quadratic effect of SOA, and the interaction of these two terms with the RSA coefficients of rule, stimulus, response and conjunction. This model was fitted only with the the random intercepts for all predictors. As reported in the main text, we found that response failures become more predictable by conjunctive representations in earlier segment of trajectories as SOA increased, *t*(1,41) = 2.85, *beta* = .004, 95% CI [.01 .05] for linear effect and *t*(1,41) = 2.85, *beta* = .004, 95% CI [.01 .05] for quadratic effect. Similar effects were observed for the rule representations, *t*(1,41) = 2.61, *beta* = .021, 95% CI [.01 .04] for linear effect and *t*(1,41) = -2.10, *beta* = -.017, 95% CI [-.03 .01] for quadratic effect, and for the response representations, *t*(1,41) = 4.20, *beta* = .036, 95% CI [.02 .05] for linear effect and *t*(1,41) = .84, *beta* = .001, 95% CI [-.01 .02] for quadratic effect.

While the longer SOA intervals led to improved behavior (Figure 3D), they did not significantly enhance the decoding of task factors in correct trials over sustained periods (Figure 4, Figure S4, Figure S5, and Table 4). Instead, the enhancement was found to be weak and transient. For example, the decoding of rules showed enhancement prior to the onset of a trigger. Additionally, when considering the results of a multilevel model that included the scores of temporal stability of linear hyperplanes of task factors (Table 5), it can be attributed that the effect of rules is due to increased separability in earlier neural trajectories, while the effect of conjunctions is related to the earlier stabilization of the hyperplane.

### Dimensionality Analysis via Simulation and Modulation of Cutoff Threshold Values

We evaluated potential limitations of the dimensionality analysis through adapted binary classification, simulating patterns of neural activities that differ in representational selectivity. Note that these patterns that are selective to low-dimensional task features or integrated task features are directly sampled from RSA models (Figure 1C). In addition, patterns of diverse conjunctions were generated by combining all task features exhaustively. These patterns could correspond to neural activities at different levels (e.g., single neurons or manifolds), and our focus is the geometric properties associated with these patterns. To accomplish this, we generated data with known dimensions (i.e., ranks) and assessed different cutoff methods (i.e., exclusive cutoff vs. overall cutoff) using two distinct simulated datasets.

First, we generated a dataset by taking a full dimensional matrix (i.e., rank = 12) for each condition (i.e., 12 unique action constellations), which were then submitted to singular value decomposition. By dropping a specific number of components (i.e., by setting corresponding singular values as 0), we generated data with variable size of dimension. Then, we added the noise with particular standard deviations and zero mean gaussian. To test the effect of different cutoff values of decoding accuracies and different level of the noise, we sampled 200 patterns and 100 trials of randomly selected conditions and submitted binary classification analysis applying the exclusive or overall cutoff method (Figure S9).

Second, we generated another dataset by simulating specific patterns of tuning properties to task factors. The pure-selective patterns were designed to be tuned to the specific condition of a particular action feature (e.g., a diagonal rule), which encompassed rule-selective, stimulus-selective, and response selective patterns. In contrast, we simulated two types of nonlinear mixed-selective patterns. The exclusive mixed-selective patterns are selective to the specific combination of action features alone (i.e., a specific action constellation). Therefore, the pure-selective patterns and exclusive mixed-selective patterns directly correspond to the unique columns of each RSA model (Figure 2C). In addition, we prepared diverse mixed-selective patterns that are uniquely tuned to every possible nonlinear mixture of task features. These diverse mixed-selective patterns correspond to the rest of binary separations that are not directly estimated by RSA (4056 = 2^12^ - 2 - (3 rule patterns + 4 stimulus patterns + 4 response patterns)) but are tested in the dimensionality analysis.

The effects of different tuning preferences of simulated patterns, the changes in cutoff threshold, and the scaling of noise in patterns on the results of dimensionality analysis were assessed by independent simulations (Figure S8; Figure S9). In each iteration of a simulation, we tested 2000 trials of randomly sampled action constellations. There were 100 patterns of specific tuning types with an independent gaussian noise of a mean of .5 and variable standard deviations. These simulated patterns were then submitted to the binary classification analysis following an identical procedure as the one for EEG. For the sake of simplicity, all patterns were simulated to produce constant activities over time. We repeated the simulation 1000 times and averaged the results over iterations.

When the cutoff threshold values for successful separations were lowered at the fixed level of noise, the overestimation of recovered dimensions (i.e., log_2_(*n_c_/T_c_*)) showed better alignment to the theoretical values of dimensionality, and avoiding an inflammatory bias introduced by lowering a cutoff threshold of decoding accuracy (Figure S8 top). The same patterns of results were observed for different levels of the noise at the fixed, low cutoff threshold (*p* = .55; Figure S8 bottom).

We further tested the selectivity of simulated patterns that are tuned for basis features of the current task in a pure-selective vs. mixed-selective (i.e., conjunctive) manner. Over the range of cutoff values and the noise levels, the estimated dimensions or the count of implementable binary separations (*n_c_/T_c_*) agreed with the expected values given the selectivity of underlying population codes (Figure S9). These simulation results confirm that the binary classification method with a low cutoff value and the exclusive cutoff method could be applied to noisy data where separability of patterns is generally low to study the geometry of neural responses.

A critical assumption of the RSA model is that there exist patterns exclusively tuned to one specific instance of the action constellation like an event-file (e.g., Figure 1A). Notably, when patterns with mixed-selectivity with this tuning were included in the population, the dimension expanded toward the maximum rank given the input conditions. We also replicated that more diverse mixed-selectivity also increases the representational dimensionality substantially, consistent with observations in the brains of non-human animals during context-dependent decisions.

Lastly, we applied a similar method of variable cutoff values to empirically observed data to test whether differences across conditions (e.g., the difference between correct vs incorrect or too late trials) are preserved (Figure 6-7). By increasing the cutoff values (from .51 to .55), we expect to decrease the probability of detecting separations as successful, which should reduce overall dimension scores. We indeed confirmed that dimension scores decrease (Supplementary Figure 10AB). However, the transient expansion of dimensions before response execution was preserved in higher cutoff threshold values. Furthermore, the functional positive relationship between conjunction and dimensionality was significantly greater compared to rule, stimulus, and response representations for ranges of cutoff values (Supplementary Figure 10C). The difference in dimension between correct and too late/incorrect trials was also preserved (Supplementary Figure 10D). Thus, this adapted binary classification with exclusive cutoff led to the same conclusions for higher cutoff threshold values, which shows robust relative differences in dimension across conditions.

### Exploratory Analysis with Rule-based Selection Task without a Response-deadline

In Experiment 1 of Kikumoto & Mayr (2020), participants engaged in a rule-based action selection task that involved three spatial translation rules, similar to the baseline task phase in the current study (Figure 2A). In comparison to the current task, the rule cue was presented 300 ms before the stimulus, and no response deadline was enforced. Participants were instructed to quickly and accurately select an appropriate response following the cue and the stimulus. Due to the absence of a requirement to extract intermediary neural states for behavioral outputs, interpreting the relationship between trial-to-trial behavior and trajectory dynamics in this task is challenging. However, even in the case of context-dependent actions initiated at the subjects’ volition, a momentary expansion of dimensionality is expected to occur before responses, thereby recruiting stable dynamics.

As an exploratory analysis, we applied a time-resolved RSA analysis, temporal generalization analysis and binary classification analysis to the response-aligned time-frequency decomposed signals using identical conditions of the analyses. Then, the dimensionality scores (*T_c_/N_c_*) were grouped into the ductile bins of strength of conjunctive representations averaged over the -800 to 0 ms time window. As shown in Figure 7, the dimension scores are strongly positively correlated with the strength of conjunctions (Figure S11). Notably, peaks of the dimensionality occurred at a similar timing of about 150 ms before the onset of responses across two studies (Figure 6 bottom and Figure S11 top). Further, these peaks occurred while optimal conjunctive hyperplanes were temporally generalizable, providing a stable readout of the conjunction of task factors (Figure S11 bottom). These findings not only replicate the main results but also underscore the significance of dimension expansion and stabilization of conjunctive subspaces.

### Stabilization of Neural Trajectories in Subspaces for Task Basis Features

To examine the impact of stable trajectories in the conjunctive subspace on action selection, we conducted RSA in combination with temporal generalization analysis (Figure 8). The resulting patterns of temporal generalization of optimal hyperplanes from one state of neural trajectories (i.e., dynamic components) to other time points (i.e., stable components) were analyzed for all basis features of the task. Figure S8 and Figure S9 display these patterns using response-aligned and stimulus-aligned signals, respectively.

Consistent with the results of multilevel models focusing on the dynamic components (Figure 5 and Figure S4), the instantaneous RSA coefficients of where the time samples for the training set and the test set match (i.e., where the black bars in each panel stand) of rule, response, and conjunctive representations were consistently higher for correct responses compared to incorrect or too-late responses.

When the decoding hyperplanes were tested on other time samples along the generalization time axis, RSA coefficients were found to be nonzero, indicating that neural trajectories are not entirely chaotically dynamic but rather exhibit overall sluggishness and stability (Figure S12 and Figure S13). The early trajectories, occurring within 300ms from the stimulus onset, displayed quicker stabilization for separating rule information compared to the trajectories associated with responses or conjunctions, which developed later (after 250ms from the stimulus onset). In contrast, the stimulus subspaces generally remained highly dynamic after the onset of stimuli and even towards the onset of responses. Furthermore, the rule, response, and conjunctive representations exhibited higher RSA scores at generalized time samples in correct response trials compared to other trials. However, this difference was not apparent for stimulus representations.

Taken together, these results indicate that the maintenance of stable trajectories for rules, responses, and conjunctions plays a crucial role in facilitating efficient and fast-paced execution of context-dependent actions. In contrast, the processing of stimuli predominantly relies on dynamic computations, which did not significantly constrain behavior in the task.

## Supplementary Tables

**Table 1.** SOA effects on RT, correct (accurate and in-time) responses, premature responses, late responses, omission errors, and commission errors.

| Phase | Effects | RT |  | Correct responses |  | Premature responses |  | Late responses |  | Omission errors |  | Comission errors |  |
| --- | --- | --- | --- | --- | --- | --- | --- | --- | --- | --- | --- | --- | --- |
|  |  | *beta* (CI) | *t*-value | *beta* (CI) | *t*-value | *beta* (CI) | *t*-value | *beta* (CI) | *t*-value | *beta* (CI) | *t*-value | *beta* (CI) | *t*-value |
| Variable SOA | SOA.linear | -.52 (-.60 -.45) | -11.74 | 5.10 (4.67 5.53) | 23.31 | 33.93 (23.56 44.31) | 33.93 | -6.48 (-6.87 -6.09) | -32.87 | -4.23 (-5.47 -3.01) | -6.73 | 3.72 (3.14 4.30) | -3.72 |
|  | SOA.quadratic | -.03 (-.07 -.01) | -3.95 | -2.35 (-2.61 -2.09) | -17.96 | -6.61 (-10.18 -3.02) | -6.61 | 1.14 (1.19 1.66) | 11.78 | .26 (.47 1.01) | 0.71 | .25 (.18 .49) | 0.25 |
| Sampled SOA | SOA.linear | -.24 (-.26 -.22) | -17.66 | 1.85 (1.71 1.98) | 27.91 | -1.54 (-1.78 -1.31) | -12.79 | 2.25 (-2.36 -2.14) | -2.26 | -1.54 (-1.78 -1.31) | -12.79 | -1.46 (-1.61 -1.31) | -18.54 |
|  | SOA.quadratic | -.08 (-.09 -.07) | -13.19 | .14 (.050 .25) | 2.93 | .40 (-.58 .23) | -4.50 | .37 (-.44 -.30) | -0.37 | -.40 (-.58 -.22) | -4.50 | -.26 (-.32 -.20) | -8.17 |

**Table 2.** Differences in RSA scores in correct vs. too-late/incorrect trials.

| Time intervals | Rule |  | Stimulus |  | Response |  | Conjunction |  |
| --- | --- | --- | --- | --- | --- | --- | --- | --- |
|  | *beta* (CI) | *t*-value | *beta* (CI) | *t*-value | *beta* (CI) | *t*-value | *beta* (CI) | *t*-value |
| Stimulus-aligned (0-500ms) | .04 (.03 -.06) | 5.58 | -.02 (.03 -.01) | -2.34 | .04 (.03 .06) | 5.20 | .03 (.01 .04) | 3.16 |
| Response-aligned (-500-0ms) | .03 (.02 .05) | 4.72 | .01 (-.02 .01) | -.68 | .05 (.03 .06) | 5.97 | .02 (.01 .04) | 3.09 |
| Trigger-aligned (-400-0ms) | .03 (.01 .04) | 3.64 | .21 (.01 .04) | 2.71 | .01 (-.01 .02) | .19 | .02 (.01 .03) | 1.99 |
| Trigger-aligned (0-300ms) | .04 (.02 .05) | 4.84 | -.05 (.02 .05) | -6.74 | .06 (.04 .08) | 7.58 | .03 (.02 .05) | 4.08 |

**Table 3.** Differences in RSA scores across short, medium, and long SOAs in correct response trials.

| Time intervals | Effects | Rule |  | Stimulus |  | Response |  | Conjunction |  |
| --- | --- | --- | --- | --- | --- | --- | --- | --- | --- |
|  |  | *beta* (CI) | *t*-value | *beta* (CI) | *t*-value | *beta* (CI) | *t*-value | *beta* (CI) | *t*-value |
| Stimulus-aligned (0-500ms) | SOA.linear | -.03 (-.04 -.01) | -3.54 | .01 (.01 .03) | 2.09 | -.01 (-.02 .04) | -1.44 | -.01 (-.02 .01) | -.38 |
|  | SOA.quadratic | .01 (-.01 -.01) | .25 | -.01 (-.01 .01) | .20 | -.01 (-.01 .01) | -.22 | -.01 (-.01 .01) | -.05 |
| Response-aligned (-500-0ms) | SOA.linear | -.03 (-.04 -.02) | -4.41 | -.03 (-.05 -.02) | -4.38 | -.01 (-.02 .01) | -.71 | .01 (-.01 .03) | 1.55 |
|  | SOA.quadratic | .05 (-.05 -.03) | .88 | -.04 (-.05 -.03) | -5.88 | -.01 (-.02 .01) | -.80 | -.01 (-.01 .03) | -.59 |
| Trigger-aligned (-400-0ms) | SOA.linear | .04 (.02 .05) | 5.35 | .14 (.13 .15) | 19.85 | .03 (.02 .05) | 4.79 | .01 (-.01 .02) | .96 |
|  | SOA.quadratic | -.01 (-.01 .01) | -.47 | -.03 (-.04 -.01) | -4.02 | .01 (-.01 .02) | 1.33 | .01 (-.01 .03) | 1.55 |
| Trigger-aligned (0-300ms) | SOA.linear | -.05 (-.06 .03) | -6.47 | -.14 (-.16 -.13) | -20.07 | .02 (.01 .03) | 2.15 | .02 (-.01 .03) | 1.80 |
|  | SOA.quadratic | -.01 (-.02 .01) | -.41 | .01 (-.01 .02) | .25 | -.03 (-.02 .01) | -.51 | -.01 (-.02 .01) | -1.43 |

**Table 4.** MLM predicting correct vs. incorrect or too late responses by the dynamic and stable subspaces, focusing on forward generalization.

| Time intervals | Effects | Rule |  | Stimulus |  | Response |  | Conjunction |  |
| --- | --- | --- | --- | --- | --- | --- | --- | --- | --- |
|  |  | *beta* (CI) | *t*-value | *beta* (CI) | *t*-value | *beta* (CI) | *t*-value | *beta* (CI) | *t*-value |
| -100-0ms | Dynamic | .04 (.02 .06) | 3.49 | .01 (-.02 .03) | .51 | .10 (.02 .13) | 7.36 | .09 (.07 .12) | 6.81 |
|  | Stable (forward) | .02 (-.01 .04) | 1.42 | .04 (.02 .06) | 3.34 | -.01 (-.01 .01) | -.55 | .05 (.02 .08) | 2.97 |
| -150-0ms | Dynamic | .05 (.02 .06) | 4.44 | .01 (-.02 .03) | .38 | .11 (.08 .13) | 7.84 | .12 (.08 .14) | 7.48 |
|  | Stable (forward) | .01 (-.01 .03) | .92 | .04 (.02 .06) | 3.81 | -.02 (-.04 .01) | -1.01 | .04 (.01 .07) | 2.58 |
| -200-0ms | Dynamic | .06 (.03 .08) | 4.56 | .01 (-.02 .03) | .5 | .11 (.09 .14) | 7.83 | .12 (.08 .14) | 7.48 |
|  | Stable (forward) | .01 (-.02 .04) | .64 | .05 (.02 .07) | 4.24 | -.02 (-.05 .01) | -1.37 | .04 (.01 .07) | 2.25 |
| -250-0ms | Dynamic | .06 (.04 .09) | 5.05 | .01 (-.02 .03) | .82 | .11 (.08 .14) | 7.38 | .12 (.08 .14) | 7.76 |
|  | Stable (forward) | .01 (-.02 .03) | .23 | .05 (.03 .08) | 4.78 | -.05 (-.05 .01) | -1.36 | .03 (.01 .02) | 2.02 |
| -300-0ms | Dynamic | .07 (.04 .09) | 5.27 | .01 (-.01 .04) | .99 | .11 (.08 .13) | 7.18 | .12 (.10 .15) | 7.64 |
|  | Stable (forward) | .01 (-.03 .04) | .04 | .06 (.04 .08) | 5.08 | -.02 (-.05 .01) | -1.40 | .03 (.01 .07) | 1.93 |
| -350-0ms | Dynamic | .07 (.05 .09) | 5.48 | .01 (-.02 .04) | .85 | .10 (.08 .13) | 7.01 | .12 (.10 .15) | 7.25 |
|  | Stable (forward) | -.01 (-.03 .03) | -.04 | .06 (.04 .09) | 5.21 | -.02 (-.05 .01) | -1.41 | .04 (.01 .07) | 2.02 |
| -400-0ms | Dynamic | .07 (.05 .09) | 5.76 | .01 (-.02 .03) | .35 | .10 (.08 .13) | 7.25 | .12 (.09 .15) | 7.02 |
|  | Stable (forward) | -.01 (-.03 .03) | -.13 | .06 (.04 .09) | 5.14 | -.02 (-.05 .01) | -1.42 | .04 (.01 .07) | 2.1 |

**Table 5.** MLM predicting correct vs. incorrect or too late responses by the dynamic and stable subspaces, including both forward and backward generalization.

| Time intervals | Effects | Rule |  | Stimulus |  | Response |  | Conjunction |  |
| --- | --- | --- | --- | --- | --- | --- | --- | --- | --- |
|  |  | *beta* (CI) | *t*-value | *beta* (CI) | *t*-value | *beta* (CI) | *t*-value | *beta* (CI) | *t*-value |
| -100-0ms | Dynamic | .03 (.01 .06) | 2.39 | -.01 (-.03 .03) | -.27 | .10 (.05 .11) | 6.27 | .08 (.05 .12) | 5.24 |
|  | Stable (both) | .02 (-.01 .06) | 1.44 | .04 (.02 .07) | 3.24 | -.01 (-.01 .01) | .47 | .06 (.03 .09) | 3.51 |
| -150-0ms | Dynamic | .05 (.02 .07) | 3.25 | -.01 (-.03 .02) | -.45 | .10 (.07 .12) | 6.68 | .09 (.06 .12) | 5.79 |
|  | Stable (both) | .02 (-.01 .05) | 1.10 | .05 (.02 .07) | 3.5 | -.01 (-.03 .03) | .13 | .06 (.02 .09) | 3.32 |
| -200-0ms | Dynamic | .06 (.03 .09) | 3.78 | -.01 (-.03 .03) | -.01 | .09 (.07 .13) | 5.86 | .10 (.07 .14) | 5.81 |
|  | Stable (both) | .01 (-.02 .05) | .77 | .05 (.03 .08) | 3.68 | .01 (-.02 .04) | .42 | .05 (.02 .09) | 3.14 |
| -250-0ms | Dynamic | .06 (.03 .09) | 3.78 | -.01 (-.03 .03) | -.01 | .09 (.07 .13) | 5.58 | .10 (.07 .14) | 5.81 |
|  | Stable (both) | .01 (-.02 .05) | .77 | .05 (.03 .08) | 3.68 | .01 (-.02 .08) | .41 | .05 (.02 .09) | 3.14 |
| -300-0ms | Dynamic | .05 (.02 .09) | 3.99 | .01 (-.02 .04) | .54 | .08 (.05 .12) | 5.57 | .10 (.06 .14) | 5.89 |
|  | Stable (both) | .01 (-.03 .04) | .47 | .05 (.02 .08) | 3.08 | .01 (-.02 .04) | .84 | .05 (.02 .09) | 2.94 |
| -350-0ms | Dynamic | .05 (.02 .09) | 3.99 | .01 (-.02 .04) | .55 | .09 (.06 .14) | 5.57 | .10 (.06 .14) | 5.86 |
|  | Stable (both) | .01 (-.03 .04) | .47 | .05 (.02 .08) | 3.08 | .01 (-.02 .04) | .84 | .05 (.02 .09) | 2.95 |
| -400-0ms | Dynamic | .07 (.04 .10) | 3.96 | .01 (-.02 .04) | .59 | .08 (.05 .11) | 5.02 | .10 (.07 .13) | 4.92 |
|  | Stable (both) | .01 (-.03 .04) | .24 | .04 (.01 .07) | 2.65 | .01 (-.02 .05) | .71 | .05 (.01 .09) | 2.60 |

## Supplementary Figures

**Figure S1.**
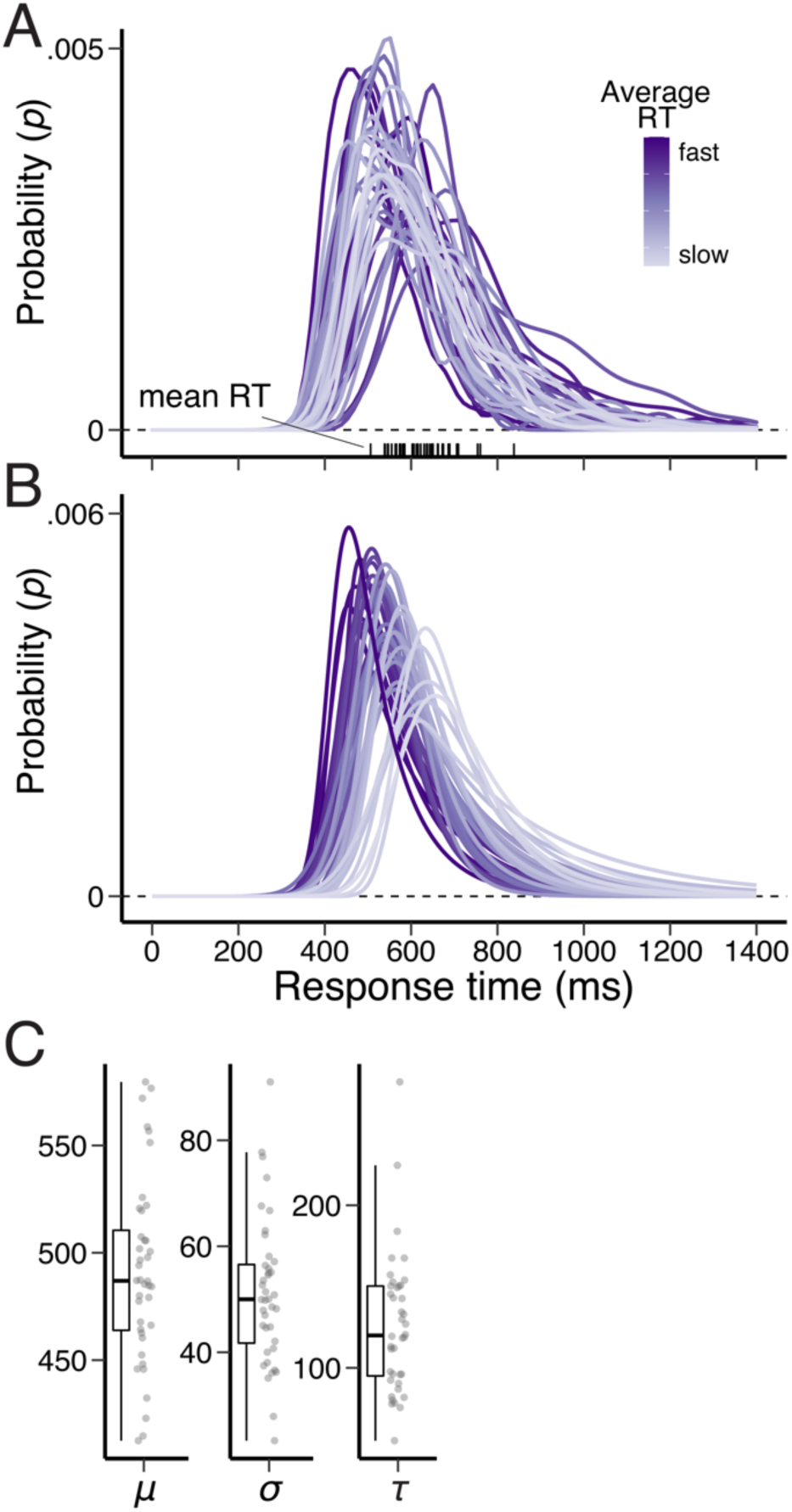
Distribution of RTs in the no deadline phase. (A) Empirically observed individual subject RT distributions in the no deadline phase of the rule-selection task. The tick marks at the bottom of each panel indicate individuals’ average RTs. (B) Individual subject predictions by a model with ex-gaussian distribution using subjects-specific estimated parameters. (C) Distributions of parameters: mean (*μ*), standard deviation(*σ*) and the skewness (*τ*) of the ex-Gaussian distribution.

**Figure S2.**
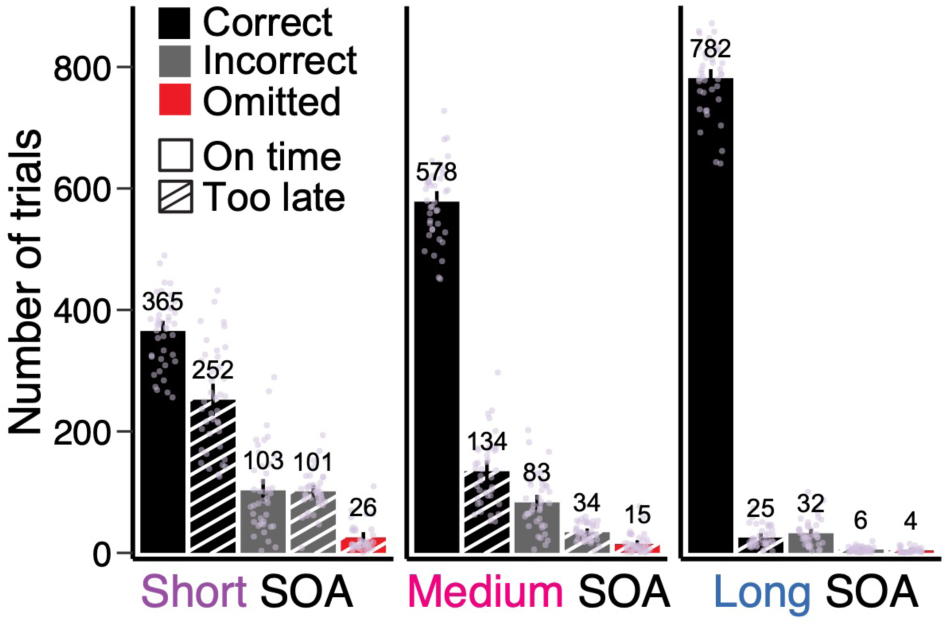
Count of different response types in each SOA condition. Empirically observed average number of trials of response errors (correct vs. incorrect) crossed by response timings (on time vs. too late) relative to subject-specific response deadlines, and response omission. Each dot represents individuals’ data points.

**Figure S3.**
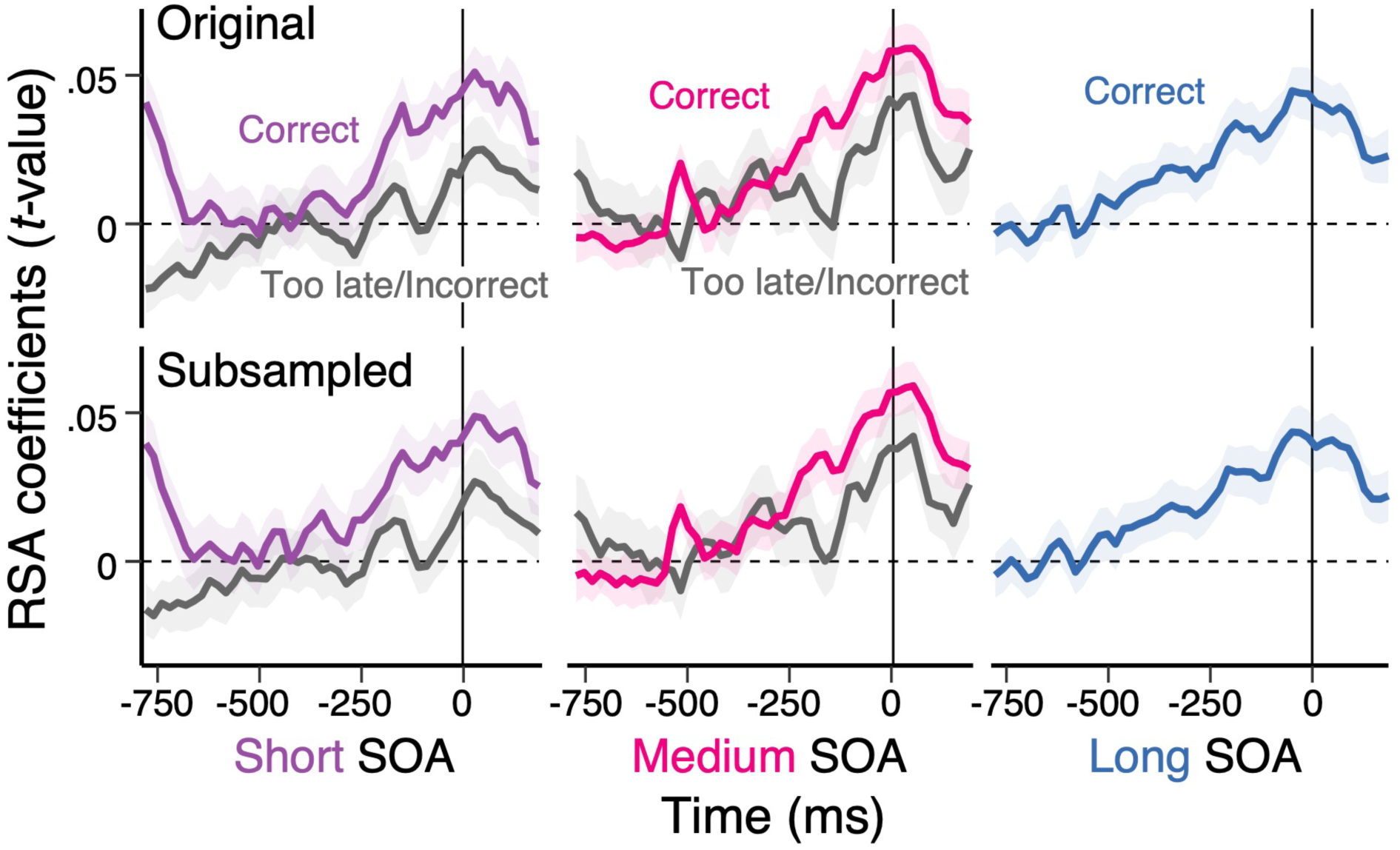
Response-aligned decoding of conjunctions using subsampling. Average, single-trial RSA coefficients (*t*-values) associated with each of the basis set task features (rule, stimulus, and response) and their conjunction that are aligned to the onset of response. The panels at the top row correspond to the conjunction in Fig.4 and the panels at the bottom row are RSA results subsampling the number of correct, incorrect and too-late responses. The left, middle, and right columns correspond to the short, medium, and long SOA conditions respectively. The two lines in each panel show the average scores in trials comparing correct vs. too late or incorrect responses. Shaded regions specify the 95% within-subject standard errors.

**Figure S4.**
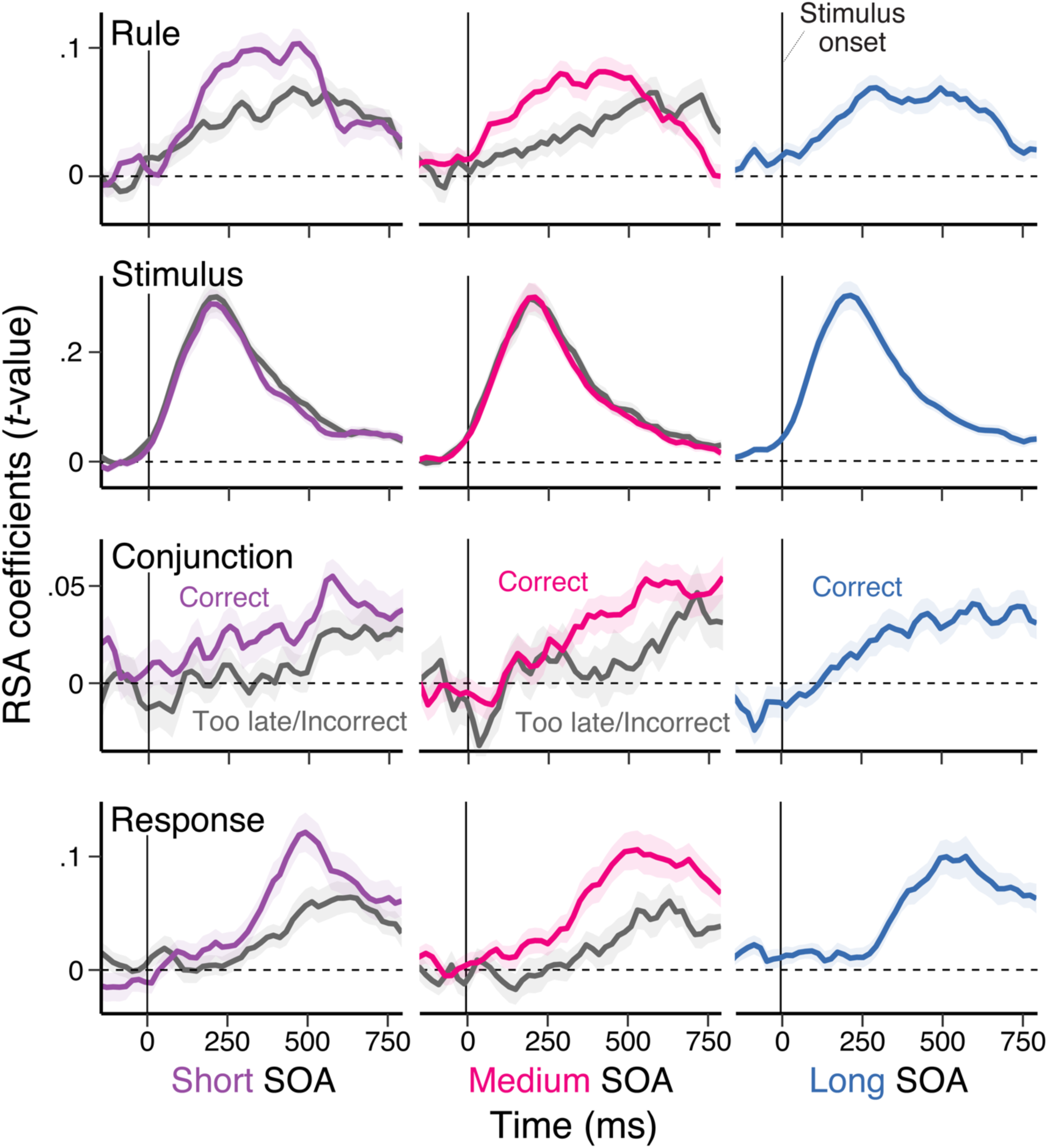
Stimulus-aligned time-course of decoding of task representations. Average, single-trial RSA coefficients (*t*-values) associated with each of the basis set task features (rule, stimulus, and response) and their conjunction that are aligned to the onset of stimulus. The left, middle and right column correspond to the short, medium, and long SOA conditions respectively. The two lines in each panel show the average scores in trials comparing correct vs. too late or incorrect responses. Shaded regions specify the 95% within-subject standard errors.

**Figure S5.**
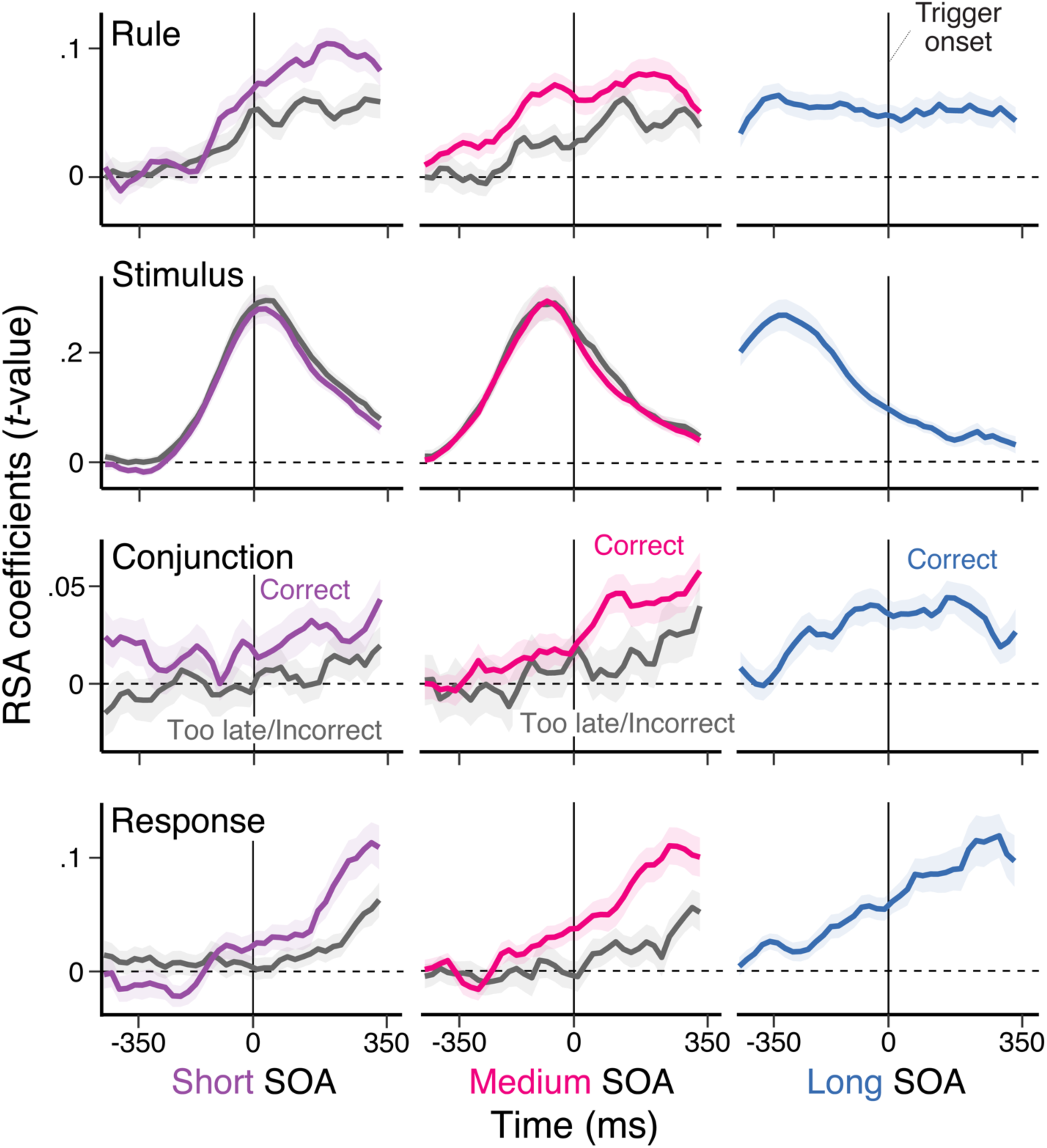
Trigger-aligned time-course of decoding of task representations. Average, single-trial RSA coefficients (*t*-values) associated with each of the basis set task features (rule, stimulus, and response) and their conjunction that are aligned to the onset of trigger for the start of response deadline. The left, middle and right column correspond to the short, medium, and long SOA conditions respectively. The two lines in each panel show the average scores in trials comparing correct vs. too late or incorrect responses. Shaded regions specify the 95% within-subject standard errors.

**Figure S6.**
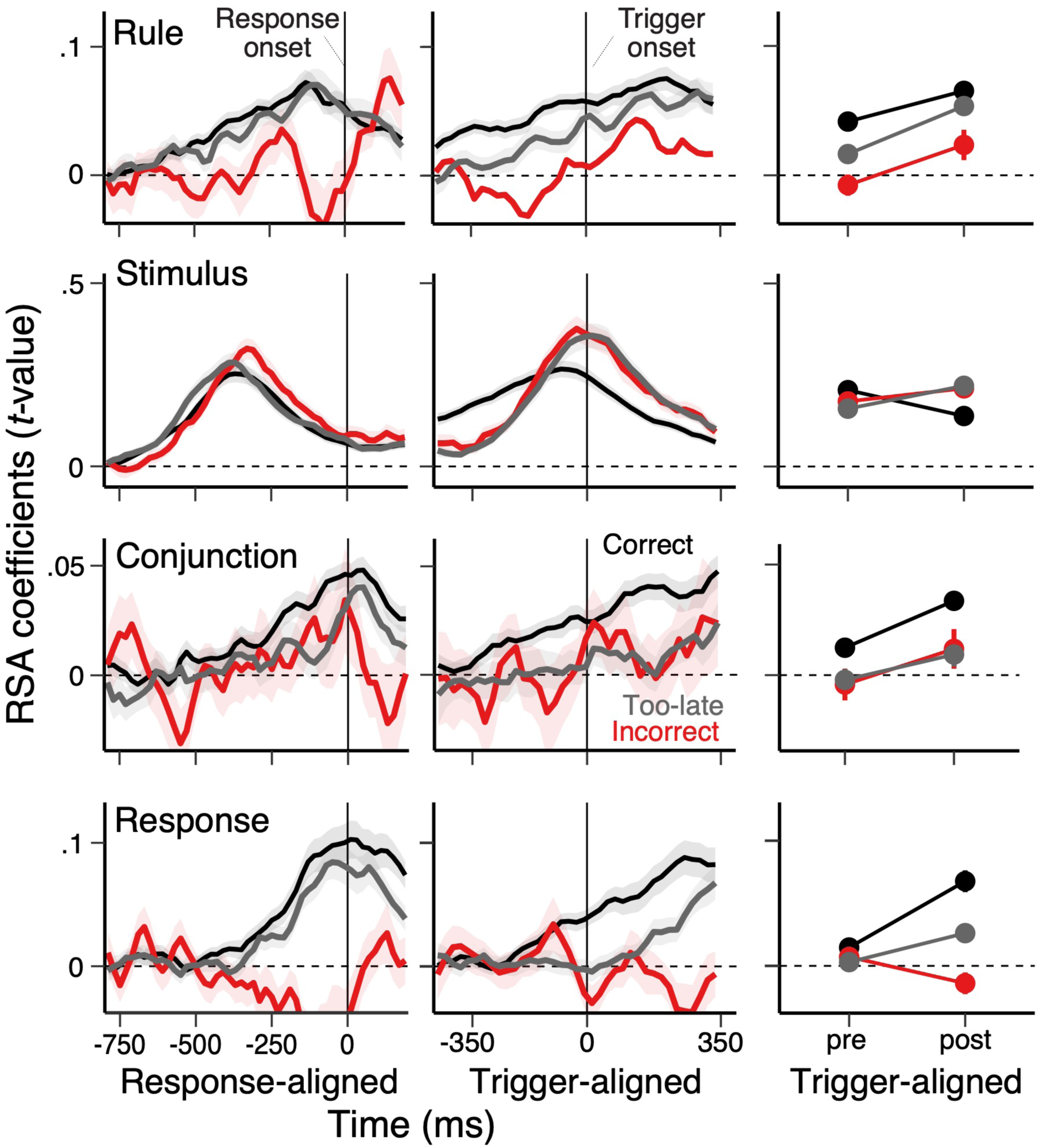
Decoding of task representations separating incorrect and too late trials. Average, single-trial RSA coefficients (*t*-values) associated with each of the basis set task features (rule, stimulus, and response) in correct, incorrect, and too late trials. The right column summarizes averaged RSA coefficients pre- and post-trigger onsets. Shaded regions specify the 95% within-subject standard errors.

**Figure S7.**
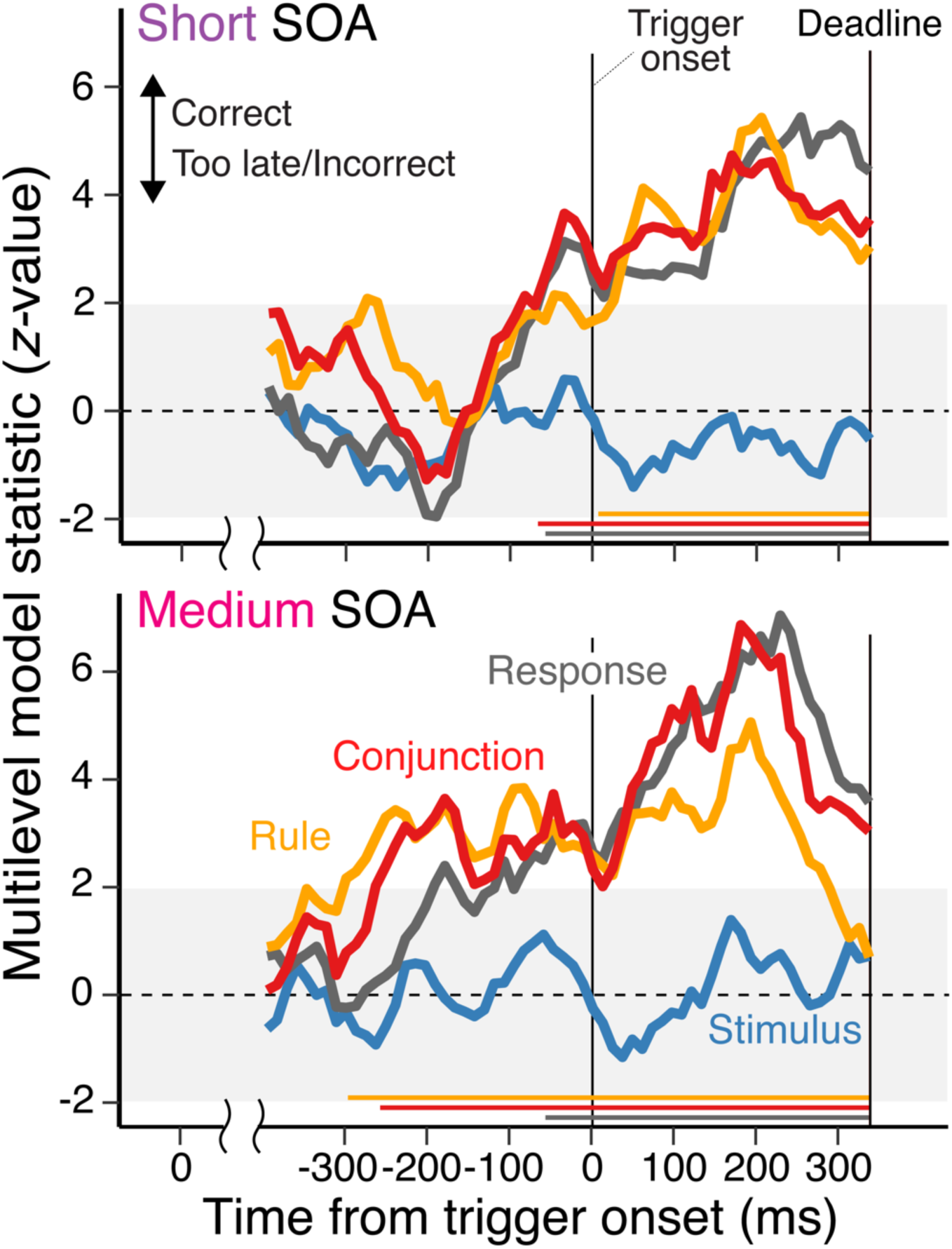
Time-course of impact of decoding of task representations on successful response selection in short and medium SOA condition. The time-course of the effect of decoded task representation strength on response selection performance. The *z*-values were obtained from multilevel, logistic regression models predicting the variability in trial-to-trial accurate and on-time responses from the strength of task representations at each moment. The signals are aligned to the onset of trial-to-trial responses. The top panel shows a result based on short SOA trials, whereas the bottom panel shows a result in medium SOA trials. In both cases, positive *z*-values indicate more successful responses as the strength of decoded representations increases. The colored straight lines at the bottom of each panel denote the significant time points using a nonparametric permutation test (cluster-forming threshold, *p* < .05, cluster-significance threshold, *p* < .05, two-tailed).

**Figure S8.**
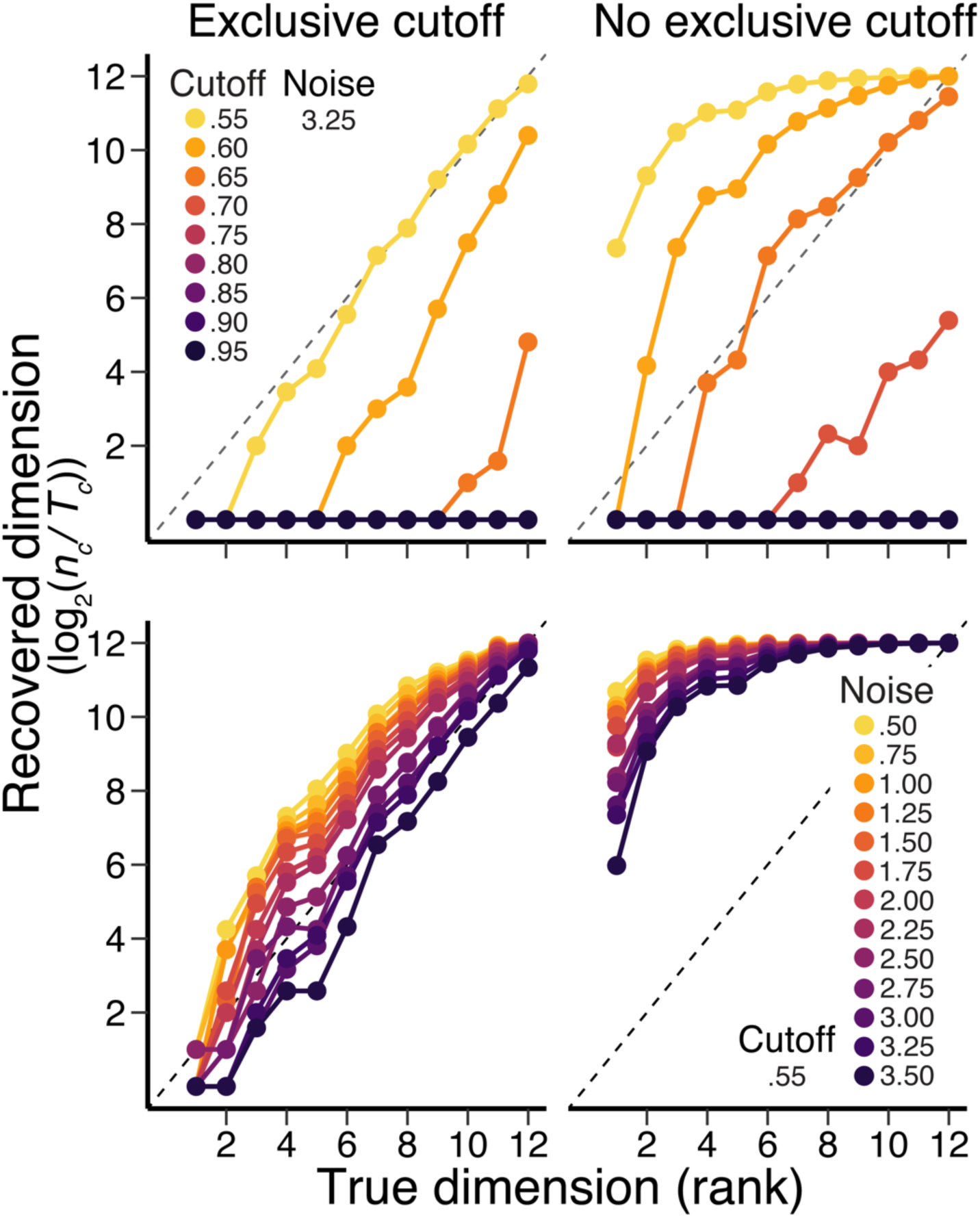
Assessment of exclusive cutoff threshold method varying the level of noise and cutoff threshold values in simulated data. The estimated dimension via the count of implementable binary classifications (separations), n_c_/T_c_, in using the simulated data with known dimension (i.e., ranks). The top two panels show the effect of changing cutoff threshold values with and without exclusive cutoff criteria on the recovered dimensionality estimates. The bottom two panels show the effects of varying the level of noise with a fixed cutoff threshold (.55).

**Figure S9.**
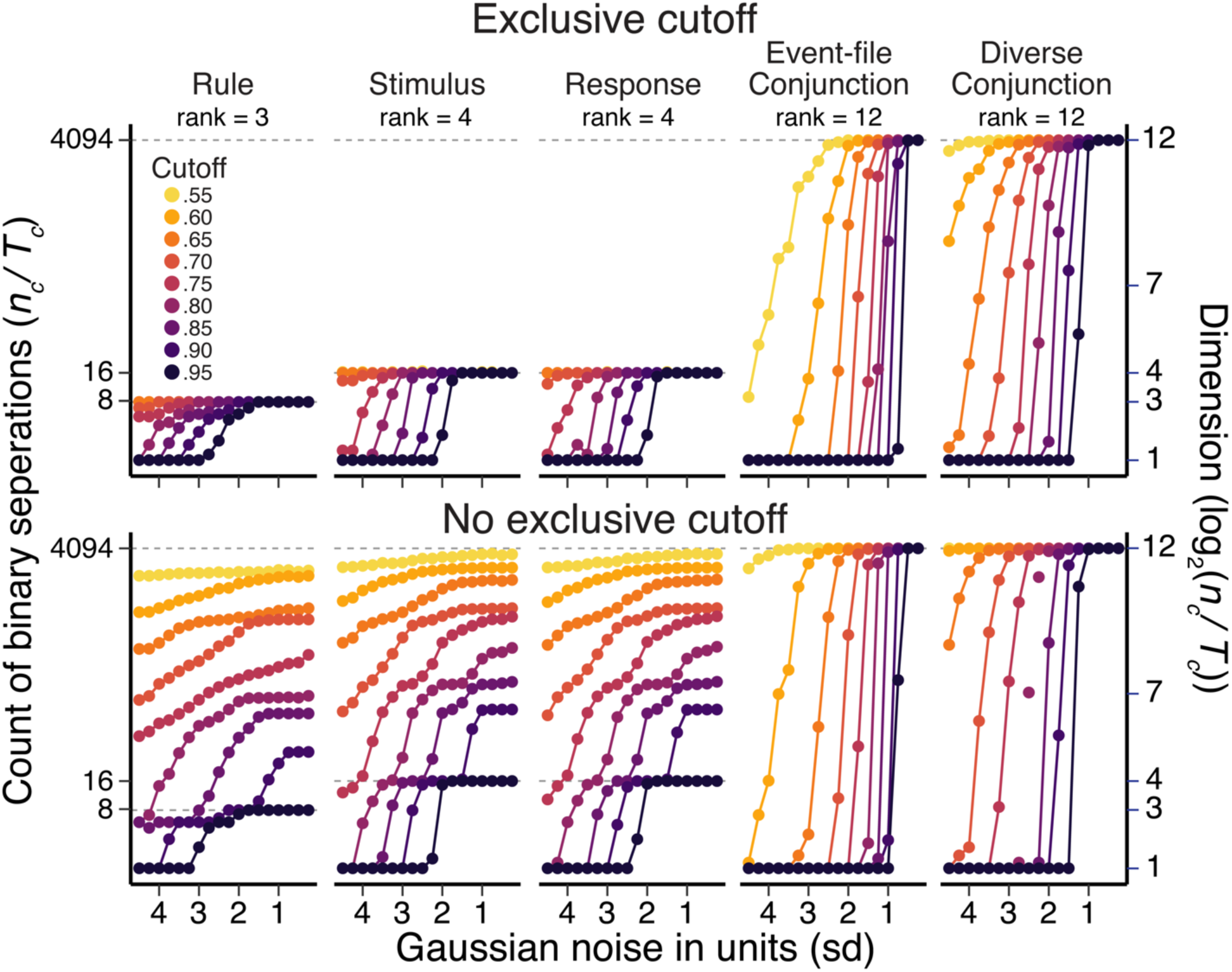
Assessment of exclusive cutoff threshold method in simulated data. The estimated dimension via the count of implementable binary classifications (separations), n_c_/T_c_, in simulated neural population responses tuned to the hypothesized task factors of the current rule-selection task. In 12 unique action constellations (input conditions), there are 4094 possible binary separations. Within each panel, *n_c_/T_c_* is computed as a function of gaussian noise and the level of cutoff threshold values where the results are summarized over multiple iterations of simulations. Each column from left side shows the result using a population tuned to specific classes of rules, stimuli, responses, exclusive combination of task factors (i.e., one-hot vector like responses or event-files) or subsample of all possible combination of task factors. The top and bottom rows show the result with or without exclusive cutoff method.

**Figure S10.**
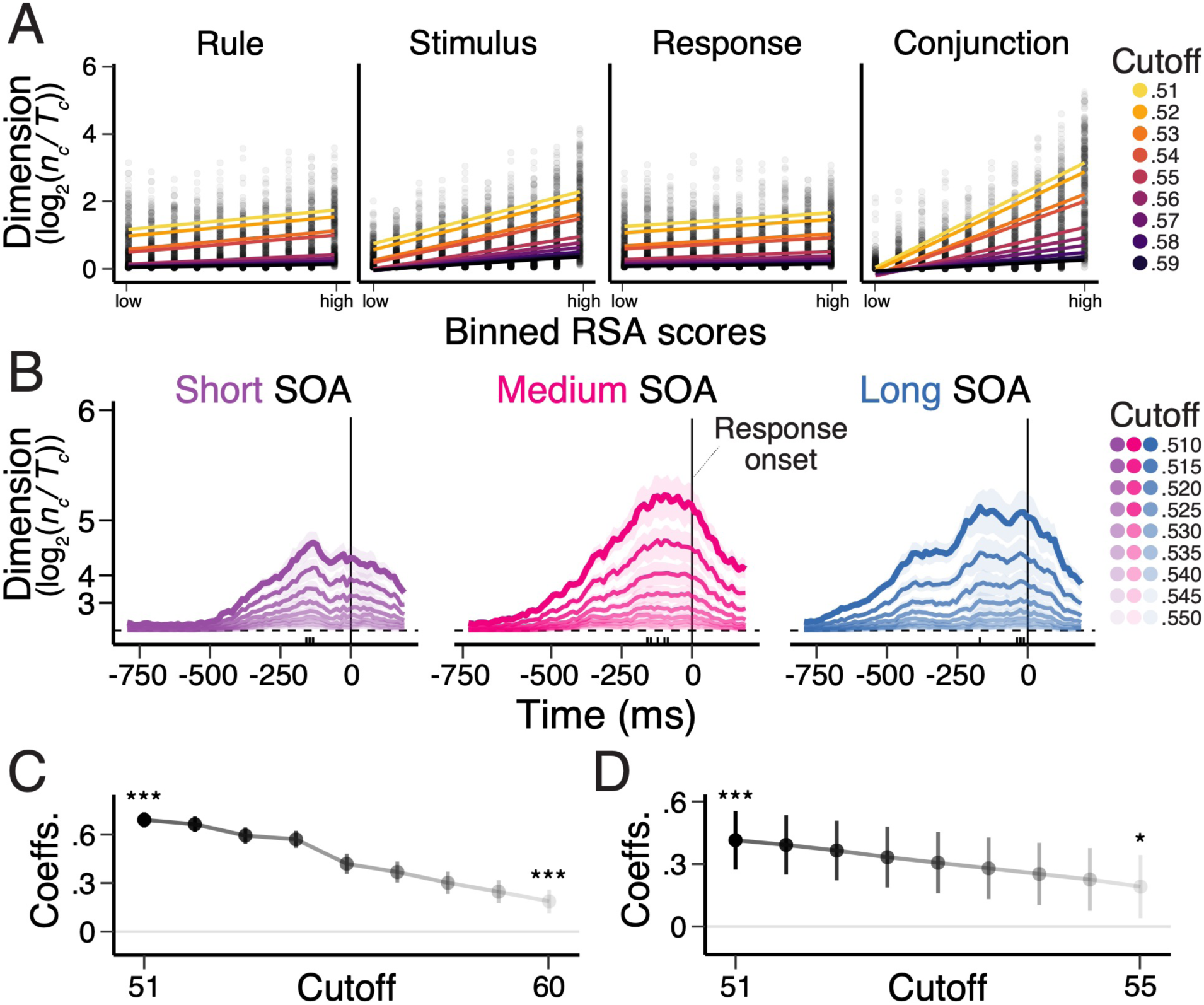
Effects of changing cutoff values of the exclusive method on estimates of task representational dimensionality. Using the identical results of binary classification in Figure 6-7, the cutoff values of the exclusive cutoff method were modulated. (A) Changes in the representational dimensionality as a function of RSA scores of each of the basis set task features (rule, stimulus, and response) and their conjunction. Different lines correspond to linear regression lines using detected binary separations with variable cutoff values. (B) Changes in dimensionality over time relative to the onset of responses. Different lines correspond to dimensions with variable cutoff values. (C) How the positive relationships in Figure 7 differ between conjunction and other task features and how they change as the cutoff value increases. (D) How the difference in correct vs. incorrect or too-late responses in dimension (Figure 6) change as the cutoff value increases, collapsing short and medium SOA conditions.

**Figure S11.**
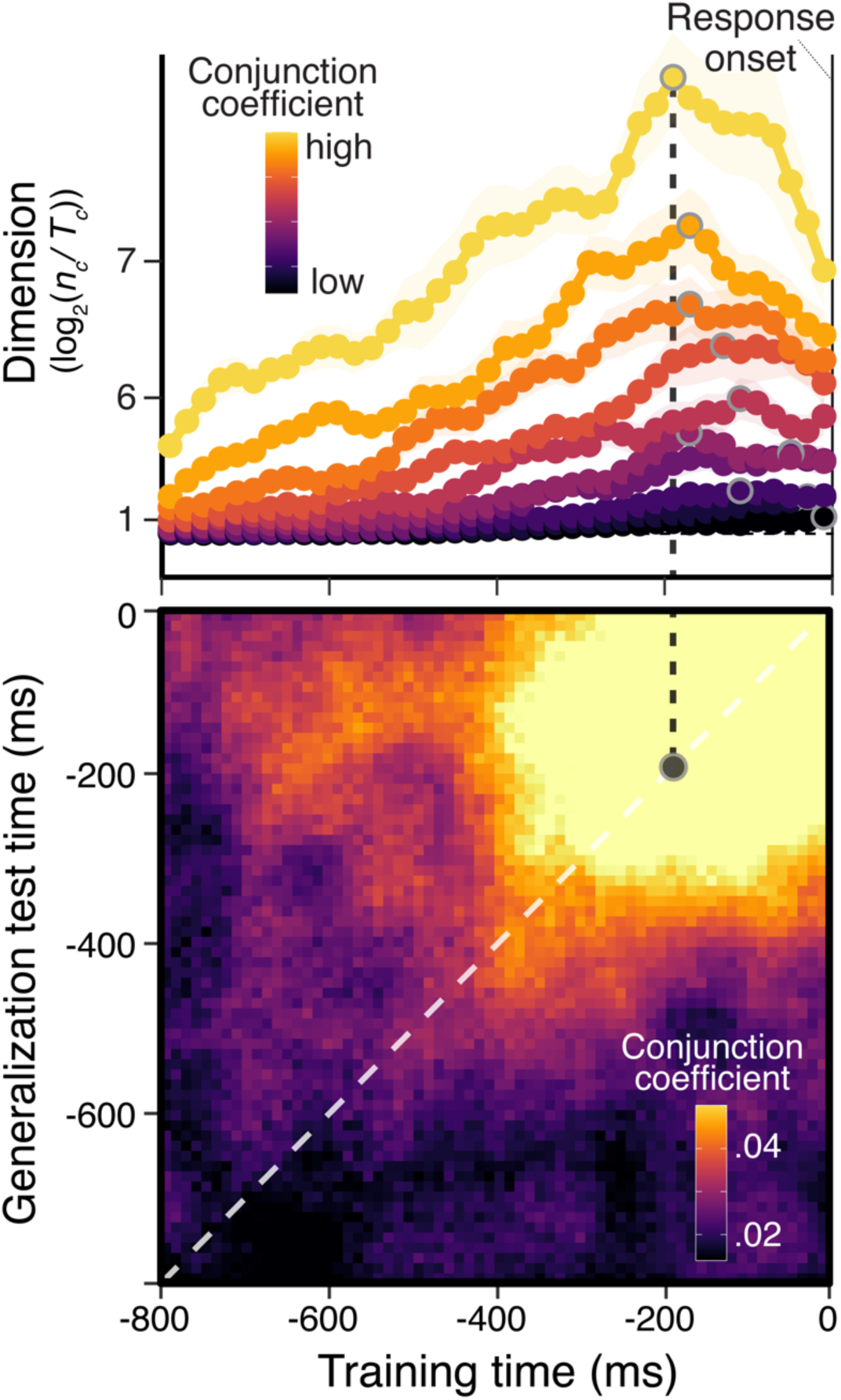
Exploratory analysis of the representational dimensionality and dynamics. Exploratory analyses of the representational dimensionality and dynamics during rule-selection task without response deadline in Kikumoto & Mayr, 2020. The top panel shows the average, dimension score during action selection. The dimensionality is estimated as the proportion of implementable binary separation, *n_c_/T_c_*, out of all possible arbitrary binary pairs. Each line corresponds to the dimension scores (*n_c_/T_c_*) calculated separately in decile bins based on the strength of conjunctive representations within subjects. The bottom panel shows the temporal generalization of linear hyperplanes that separates conjunctions summarized as the decoding matrix.

**Figure S12.**
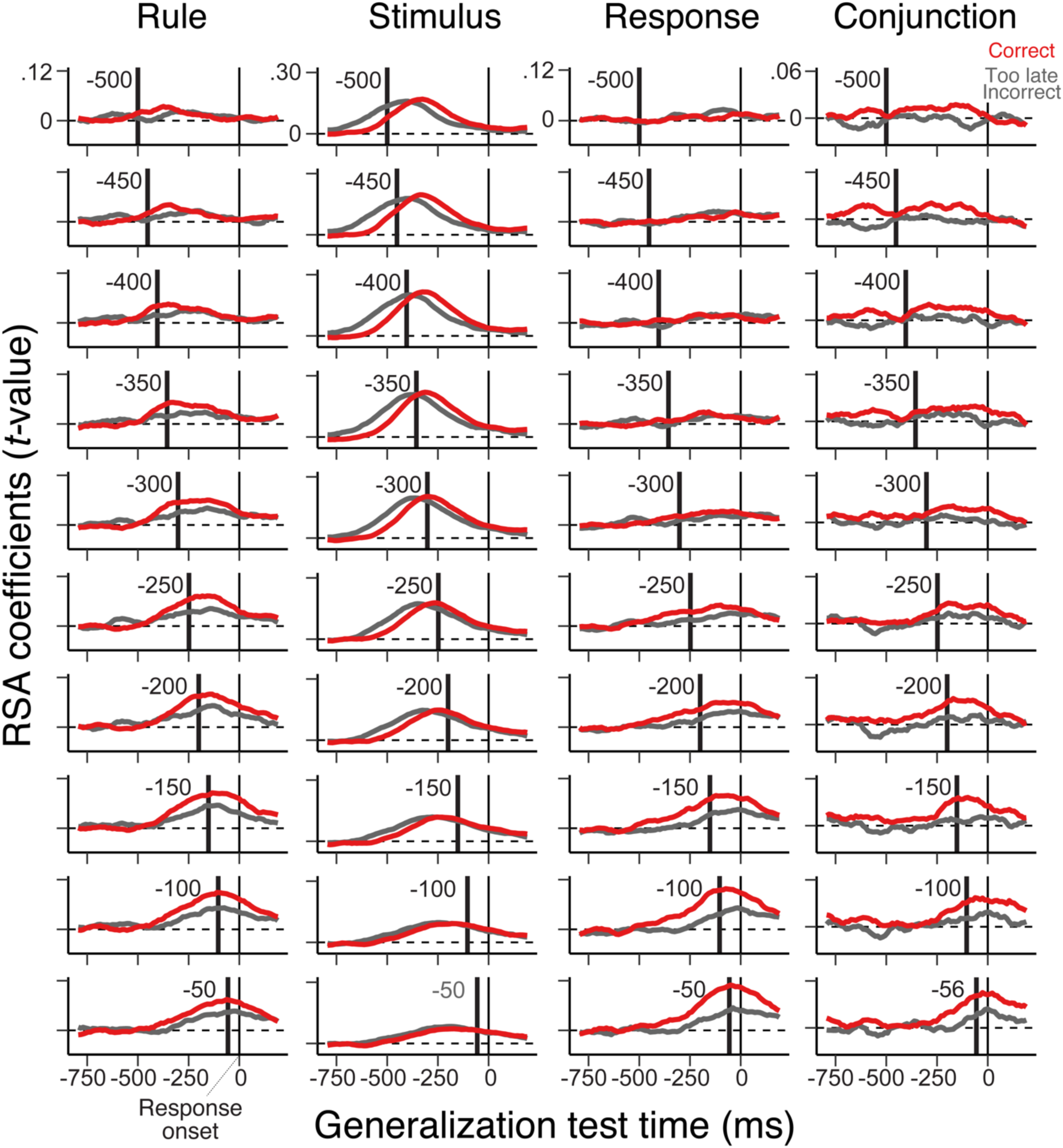
Stability of the representational dynamics of rule, stimulus and response subspaces using response-aligned signals. Temporal generalization of linear hyperplanes that separate basis task factors and the conjunction. Each column shows the results for different task factors. Within each panel, the black bar denotes the center of 50 ms time windows (±20 ms relative to the center) of signals to obtain hyperplanes of decoders (i.e., dynamic components). The trajectories show the results of temporal generalization to other time samples (i.e., stable components), separately for correct vs. incorrect or too-late responses. The 0 ms point on x-axis denotes the onset of responses.

**Figure S13.**
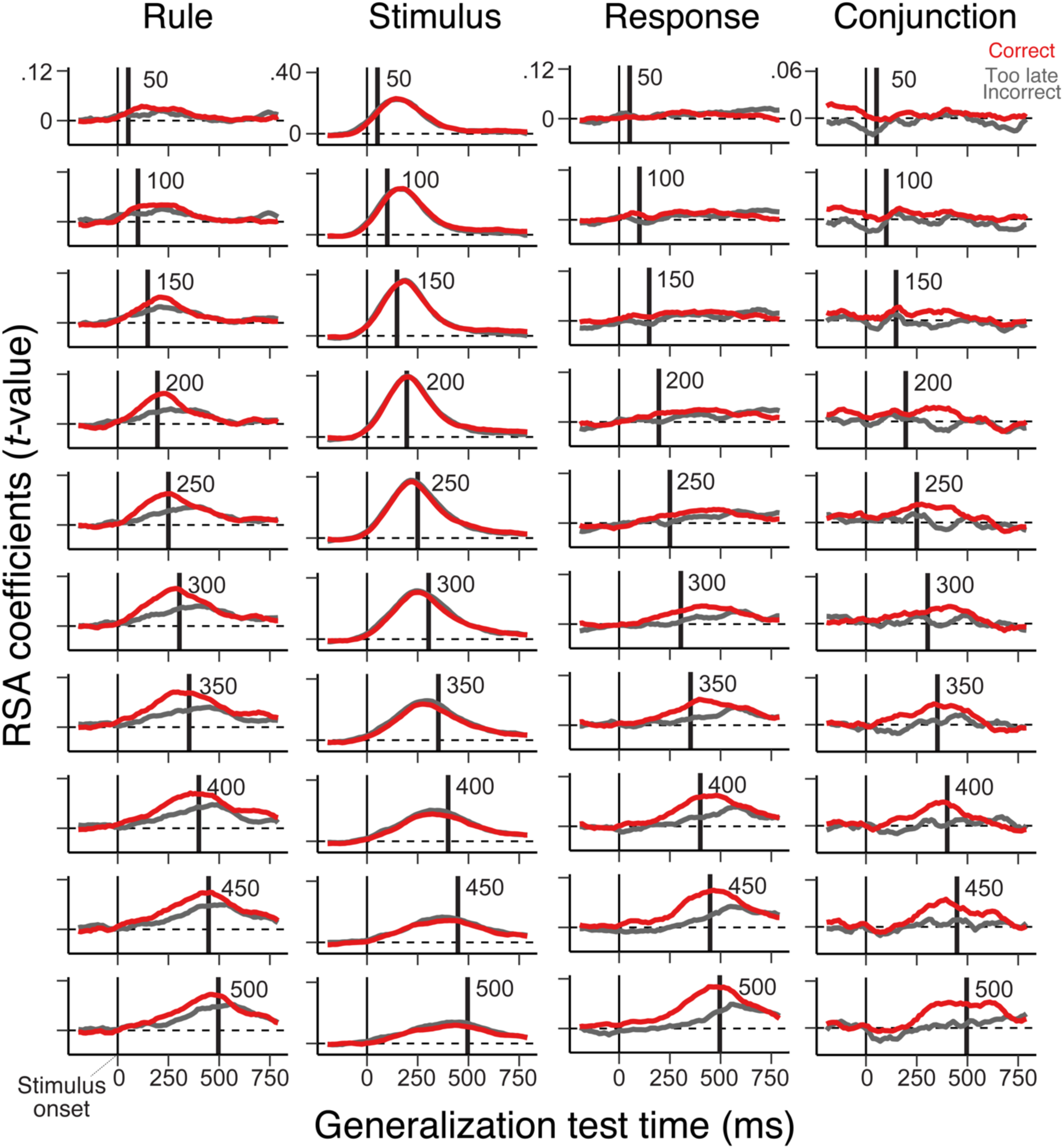
Stability of the representational dynamics of rule, stimulus and response subspaces using stimulus-aligned signals. Temporal generalization of linear hyperplanes that separate basis task factors and the conjunction. Each column shows the results for different task factors. Within each panel, the black bar denotes the center of 50 ms time windows (±20 ms relative to the center) of signals to obtain hyperplanes of decoders (i.e., dynamic components). The trajectories show the results of temporal generalization to other time samples (i.e., stable components), separately for correct vs. incorrect or too-late responses. The 0 ms point on x-axis denotes the onset of stimuli.

## Notes

https://neurodata.riken.jp/

### Competing Interest Statement

The authors have declared no competing interest.

### Summary of Updates

We performed more control analyses related to the dimensionality analysis (changing cutoff values) and single-trial RSA (adjusting the number of trials), and added the summary of each response type.

